# Routing of task-relevant information in mouse PPC during continuous visuomotor control

**DOI:** 10.64898/2025.12.22.696069

**Authors:** Poojya Ravishankar, Harrison A. Grier, Matthew T. Kaufman

## Abstract

Posterior Parietal Cortex (PPC) exhibits tuning to many variables, including strong representations of visual information, movement, and behavioral biases. Whether PPC communicates all these variables to other areas is less clear. We examined PPC activity in mice performing a novel, closed-loop, 2D visuomotor joystick task that required animals to act exclusively on a task-relevant axis of visual motion. To determine what components of PPC’s representation were sent to M1, we performed two-photon calcium imaging of layer 2/3 neurons in contralateral PPC of expert mice with PPC-M1 projection neurons identified via retrograde tracing. Consistent with previous results, PPC neurons exhibited random mixed selectivity and were typically most strongly modulated by joystick movement. Most of the visually responsive neurons were more strongly modulated by task-relevant than task-irrelevant visual motion. Encoding in labeled PPC-M1 neurons was similar to encoding in unlabeled neurons, with one major exception: unlike the task-relevant visual enrichment in unlabeled PPC neurons, task-relevant and task-irrelevant visual motion were encoded at similarly weak levels in PPC-M1 neurons. This argues that although PPC encodes a mix of visual, movement and other information, the PPC-M1 pathway is dominated by movement information and does not propagate PPC’s learned enrichment of task-relevant visual signals.

## Introduction

While decision and movement are intimately linked, it remains unknown how abstract decisions are converted into detailed movement commands. Large-scale recordings have demonstrated that information about numerous variables, including both decision and movement^1–6^, are widely distributed across cortex. Even in the secondary visual areas, often thought of as a largely-feedforward hierarchy^7^, some neurons within a single area may respond to low-level features of a stimulus (e.g., oriented edges) while others respond to high-level features (e.g., the animal’s position on a track)^8^, and information about movement is strongly present throughout visual cortex^3,5,9^. Nonetheless, sensory information enters only in some cortical areas and movement commands are sent from others. We therefore ask a fundamental question about the structure of computation across cortex: how is the directional flow of neural processing achieved?

To disentangle feedforward from feedback signals in a brain area, the activity of neurons projecting to a downstream (or upstream) target can be compared to other neurons in the area. Prior work has argued for three different hypotheses. First, connections might follow the ‘linkage principle’, where the upstream area transmits the signals that most resemble the activity of its target^10,11^. Alternatively, connections might instead avoid sending target-like signals, to prevent ‘reverberant’ positive feedback loops^12,13^. Finally, connections might perform ‘complete transmission’ of the area’s representation.

Evidence exists for each hypothesis under different circumstances. The linkage principle is perhaps the natural hypothesis from considering feedforward networks, and two empirical results from early sensory areas in mouse support it. First, V1’s axon terminals in extrastriate areas resemble their targets, though that refinement is not present in the output neurons of V1^14^; and second, S1-to-S2 projection neurons preferentially carry sensory signals about object properties, while S1-to-M1 projection neurons preferentially carry information about object presence^15^. The reverberation-avoidance hypothesis is supported by recordings from mouse Posterior Parietal Cortex (PPC), a visuomotor decision area^16^. Mostly-visual signals were present in PPC-M1 neurons and axon terminals during a voluntary eye movement paradigm^13^ and visual signals were enriched over bias variables in the PPC-M2 pathway in a visually-informed lever-pull^12^. Finally, the complete-transmission hypothesis is supported by indirect evidence from PPC to motor areas, where motor areas received both relevant and irrelevant visual information during flexible visual tasks in both rats^17^ and monkeys^18,19^.

PPC is a particularly interesting target for understanding signal routing in rodents, because it offers the possibility to understand routing at two levels: routing of overall classes of information, such as transmitting mostly visual vs. motor information; and routing based on learned task-relevance. PPC is causally involved in visuomotor tasks^16,20–22^, with a connection to motor cortices specifically implicated in a joystick task^12^, and it is more active during visuomotor task performance than during passive viewing of stimuli^20^. PPC has responses that mix visual and motor^8,23–25^ and contain information about trial history^26,27^. Finally, PPC responds in a graded fashion to stimuli instead of exhibiting a categorical response, unlike motor areas^28^, but connects strongly and directly to the forelimb area of M1 as well as to the secondary motor area^7,29–32^. Thus, the pathway from PPC to M1 is a “hotline” from higher-order vision to the motor periphery.

Here, we ask how computations underlying visuomotor decision-making proceed from PPC to M1. We trained mice to perform a new continuous-control joystick task informed by a drifting visual stimulus, where one axis of motion (e.g., the vertical axis) is always relevant and the other is always irrelevant. We used two-photon calcium imaging to record the activity of over 12,000 PPC neurons with the PPC-M1 projection neurons identified, then fit encoding models to understand what information was present in each neuron. Consistent with previous results^33,8,24,34,23^, PPC exhibited mixed selectivity for a large number of task variables. We then investigated signal routing at the two levels described above. At the signal-class level, we found that the PPC-M1 projection acts consistently with the linkage principle: PPC-M1 neurons were more purely motor-encoding than other PPC neurons. At the learned-task level, we found that task-relevant visual information was strongly enriched in the overall population relative to task-irrelevant information, but that surprisingly this was not the case in the PPC-M1 neurons. Together, these results suggest that selective routing of motor information over visual is present in the PPC-M1 pathway, but that selective routing of task-relevant vs. irrelevant sensory information is not – despite a lifetime of training with a consistent contingency.

## Results

### Novel visuomotor paradigm to probe selection of task-relevant information

To understand how behaviorally relevant information is structured and propagated from PPC to motor cortex during complex visuomotor control, we required a behavioral task with two key features. First, we needed to engage the link between visual processing and movement instruction as strongly as possible. Second, behaviorally relevant and -irrelevant visual information needed to be presented on an equal footing. This latter feature provided several benefits. First, it ensured that selection of task information was purely a learned visual-motor association. Second, it permitted training some animals with one dimension of the stimulus relevant and other animals with the other dimension relevant, to validate that any differences were not innate.

In our “TextureDrift” task, the mouse first held a paw rest with one paw and a two-axis joystick with the other, then was presented with a random, splotchy, black-and-white visual texture^35^ drifting diagonally (Fig. 1a). The mouse was trained that only one dimension of the stimulus – vertical for some mice (A1-A3), horizontal for others (A4, A5) – was task-relevant. After a 250ms observation-only period in which the joystick was movable but uncoupled from any outcomes, the mouse could alter the relevant axis of the stimulus’s drift by moving the joystick (Fig. 1d-f). Because the position of the joystick mapped to adding an instantaneous velocity vector to the drifting stimulus, the drift velocity implicitly specified a target location for the joystick on each trial. Reaching the target required movements of 3.2-6 mm, though movements were typically 8-12 mm. Crucially, only one dimension of movement of the joystick affected the drift of the stimulus. The mouse was rewarded for stopping or reversing the motion of the stimulus on the relevant axis for at least 100ms or failed the trial if it released the joystick or moved the joystick more than very briefly in the opposite direction by the same distance that would have garnered a reward (into the “failure zone”; Methods). Importantly, which axis was task-relevant was fixed throughout the mouse’s entire life. Mice reached stable performance after 2-3 months of training (see Methods, Supp. Fig. 1), and performed large numbers of trials: 421 per session on average. An average of 82% of trials were rewarded (60-95% across mice, Supp. Fig. 2), but we restricted analysis to “clean” trials where the animal never moved the joystick into the failure zone (60-70% of all trials were clean successes, 66-85% for easier trials; range across mice; Fig. 1b).

**Figure 1.**
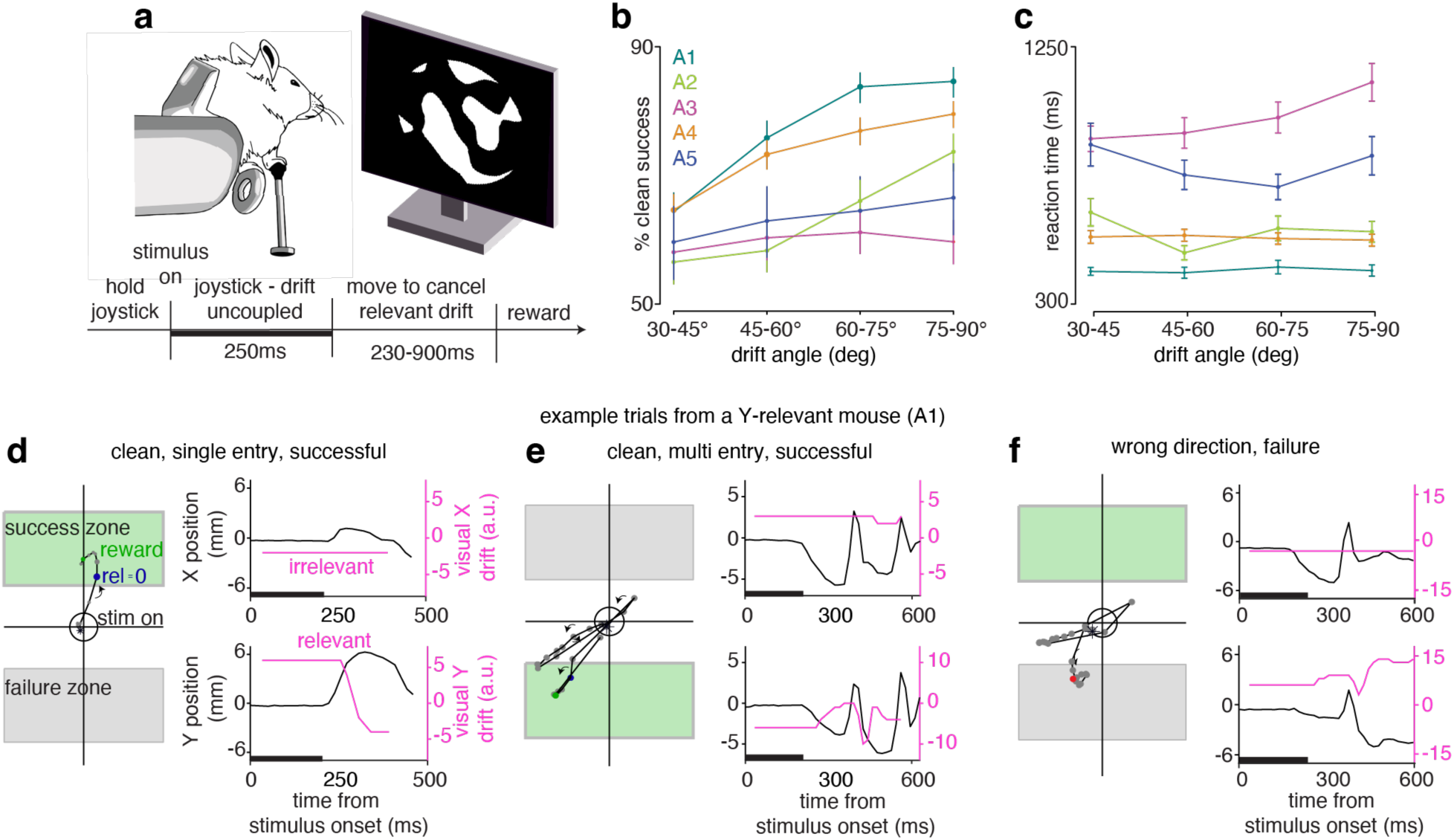
A novel closed-loop visuomotor control task with task-relevant and irrelevant visual information. (a) Schematic of the task setup (top) and trial structure (bottom). (b) Percentage of trials that were successful and had “clean” movements as a function of visual drift angle. (c) Reaction time (Methods) as a function of visual drift angle. (d) Left: Joystick position across time during a simple, clean trial depicted on the task-workspace; blue and green dots denote time of target entry (relevant drift = 0) and reward delivery. Right: instantaneous joystick position (purple) and visual drift (red) decomposed into X (dotted, irrelevant) and Y-axis (solid, relevant) across time for the trial shown to the left. Black rectangle denotes the decoupled time period (0-250ms after stimulus onset). (e) Same as (d), for a trial with multiple entries into the target zone. (f) Joystick position during an example trial when it was moved in the wrong direction/failure zone.

**Figure 2.**
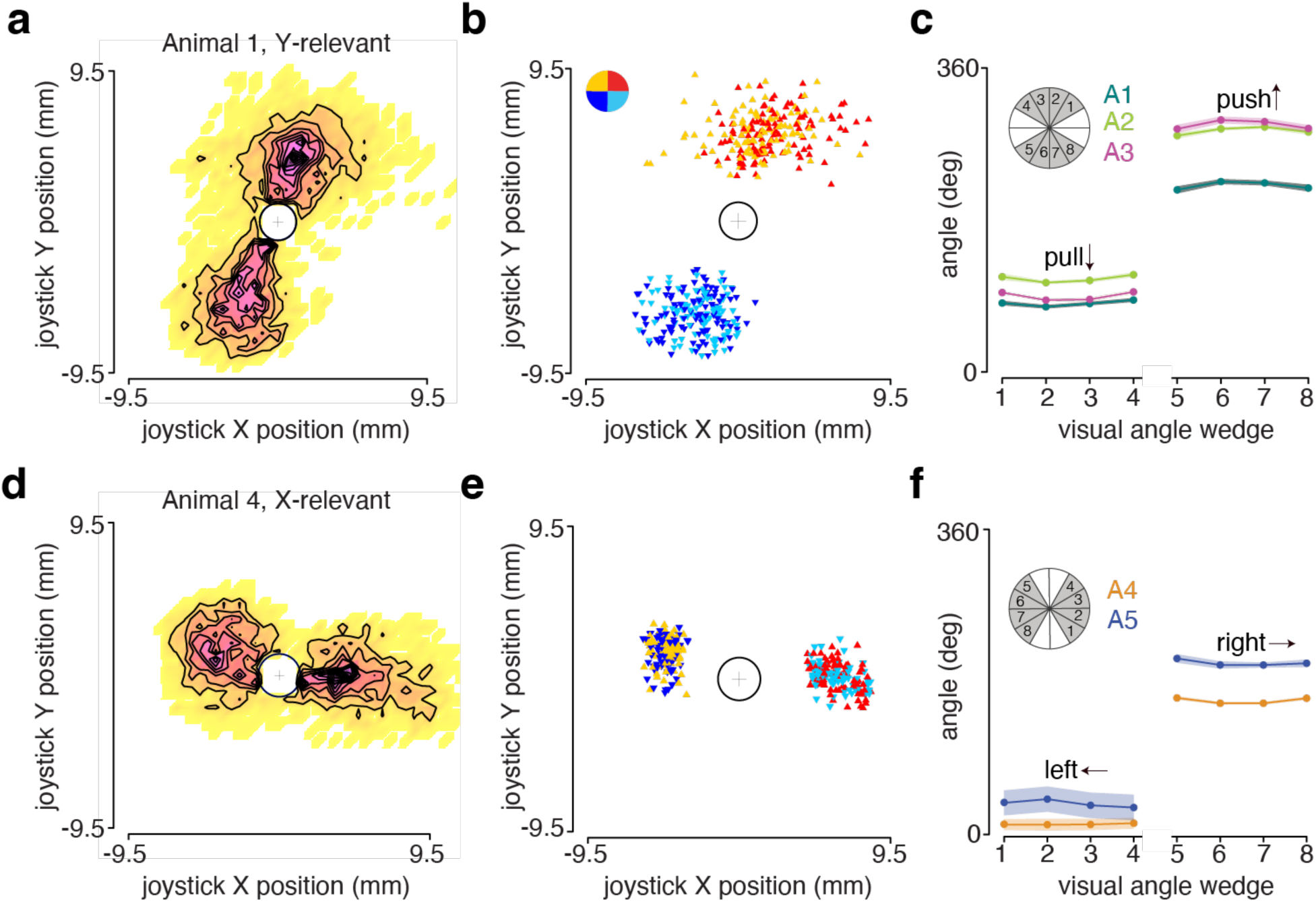
Mice successfully learn to act only on task-relevant visual information. (a) Binned joystick position (with contours) in a session from a Y-relevant mouse (A1). Central fixation zone (white circle) excluded. (b) Joystick position at the outcome sample for different trials, colored based on the quadrant of the visual drift. Inset color wheel shows quadrant color key opposite to the visual drift vector. Same mouse as in (a). (c) Joystick movement angle as a function of visual direction for all Y-relevant mice (n=3). Inset: visual direction grouping schematic. Gray indicates the wedges from which visual directions were sampled. White denotes directions that were not sampled in the task. (d) Same as (a) but for an X-relevant mouse (A4). (e) Same as (b) but for the mouse shown in (d). (f) Same as (c) for all X-relevant mice (n=2).

We first verified that expert mice learned to perform information selection by acting on the relevant axis of the visual drift and ignoring the irrelevant axis of the drift. Joystick movement was much greater in the relevant axis (Fig. 2a,d; example sessions for other mice in Supp. Fig. 3a-c, top panels), and the average angle of the joystick movement did not depend on the irrelevant drift (Fig. 2c,f). This equal average angle did not reflect idiosyncratic distributions: single-trial movement endpoints similarly did not reflect the direction of the task-irrelevant drift (Fig. 2b,e; Supp. Fig. 3a-c, bottom panels). Moreover, task-relevant visual information was the strongest single-trial predictor of movement direction in a logistic regression model with task-relevant visual information and strategy variables (Supp. Fig. 4a-e). Although stronger relevant information increased accuracy across all animals (Fig. 1b), the effects on reaction times (RTs) were less consistent, potentially hinting at different strategies (Fig. 1c). However, accuracy was higher on trials where mice waited at least 200-300ms to respond (Supp. Fig. 4g), suggesting that a minimum integration time improved performance.

**Figure 3.**
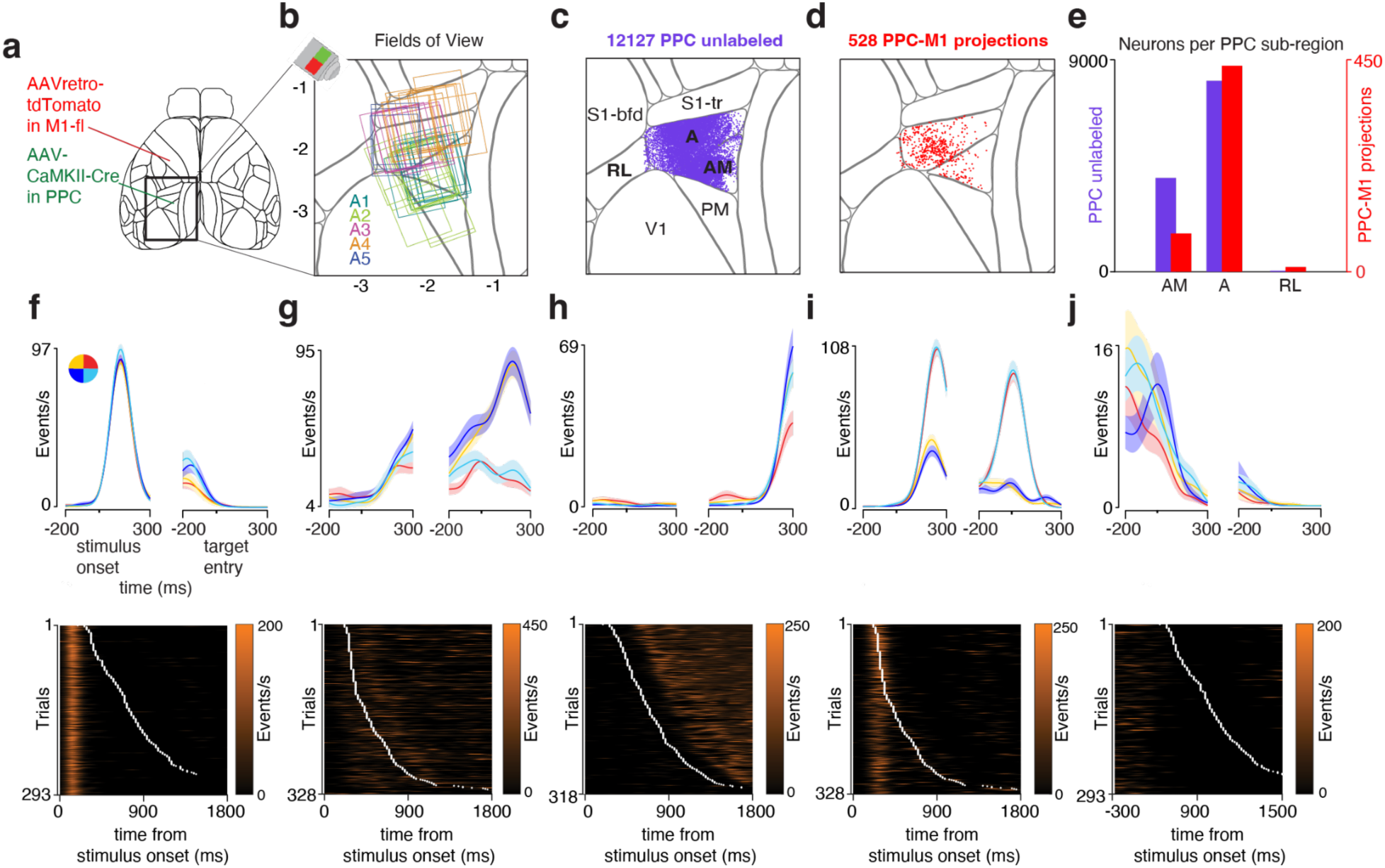
Experimental strategy and PPC responses during closed-loop forelimb control. (a) Schematic of the Allen CCF with the experimental strategy. AAV-CaMKII-Cre was injected in PPC and adjacent visual areas to cause GCaMP6f expression. AAVretro-CAG-tdTomato was injected in forelimb M1 (M1-fl) to retrogradely label PPC-M1 neurons with tdTomato. (b) Two-photon imaging fields of view (FOVs) from all mice superimposed on the Allen CCF. (c) Locations of imaged neurons that did not have red labeling (“unlabeled”). (d) Locations of red-labeled M1 projection neurons. (e) Bar plot showing the total number of unlabeled and M1 projection neurons within each constituent area of PPC. Note different axis scalings. (f-j). Top: peri-event time histograms (PETHs) of example neurons during the task, with activity aligned to the stimulus onset and to target entry. Traces correspond to averaged and smoothed responses for each visual quadrant; each plot shows one neuron. Shaded regions represent the SEM. Bottom: raster plots for the same neurons. White ticks represent the time of target entry in each trial. Neurons were chosen to represent the diversity of responses.

**Figure 4.**
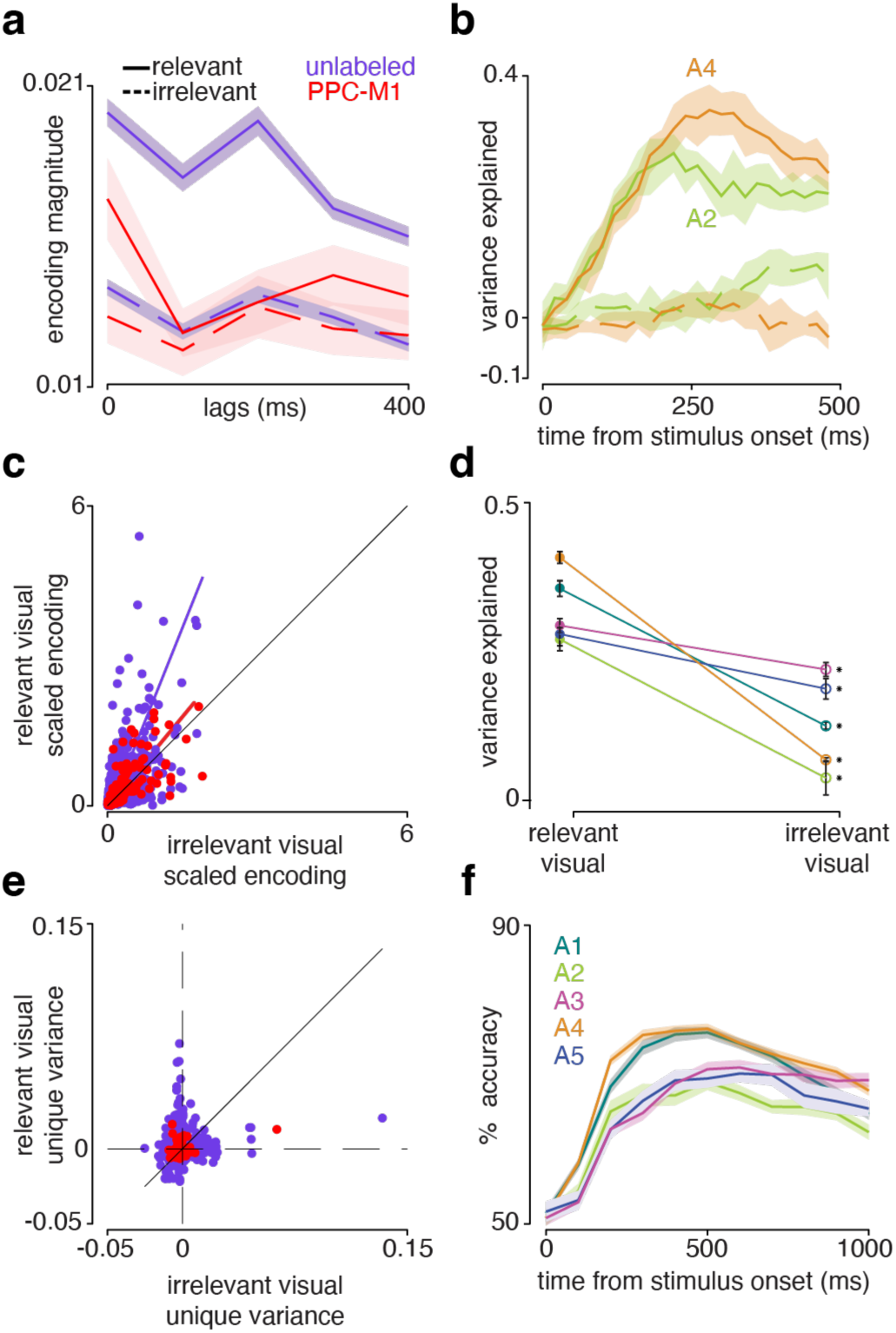
Encoding and decoding of task-relevant and irrelevant visual drift in PPC. (a) Population-averaged task-relevant (solid) and irrelevant (dashed) visual encoding kernels for unlabeled PPC (purple, n=1786) and PPC-M1 (red, n=120) neurons. Shaded regions represent the SEM. (b) Cross-validated variance explained by the instantaneous decoder for task-relevant (solid) and task-irrelevant (dashed) visual information across time for two example animals. All fitted neurons. SEMs are across sessions. Two example animals shown. (c) Single neuron encoding magnitude for the relevant and irrelevant visual variables (quadratic mean across lags) for unlabeled PPC and PPC-M1 neurons along with the total least-squares fit lines. The encoding magnitude of each neuron is scaled by the standard deviation of its firing rates (see Methods). (d) Paired plot comparing the cross-validated variance explained by the instantaneous decoder for task-relevant (solid) and irrelevant (open) visual information. Each line is a different mouse. Error bars are across sessions. All fitted neurons. (e) Unique variance captured by the task-relevant and irrelevant visual variables in unlabeled (purple, n=1932) and PPC-M1 (red, n=128) neurons. (f) Cross-validated accuracy across time for a decoder trained to categorize the direction of task-relevant visual information. Each line is a different mouse. Shaded regions represent the SEM across sessions.

Performing this task was challenging for the mice, and they produced a variety of joystick movements across trials. Many trials elicited simple movements, where the mouse went directly to the target zone and held it long enough for a reward (Fig. 1d). Other trials required multiple attempts to hold the joystick in the target zone for long enough (Fig. 1e). Excursions into the opposite zone caused failure (Fig. 1f). For the first three mice (A1-A3), brief excursions into the opposite zone did not cause failure. To ensure that the animals were not performing above chance due to a degenerate strategy, for behavioral analysis (Fig. 1b,c) we excluded trials in which they moved the joystick into the failure zone at any point.

Disrupting PPC activity alters behavior in a variety of previously-studied visually-guided behaviors in rodents, though specifics of the behavioral effects vary^16,20–22,26–28^. As a simple first verification that PPC was causally involved in our task, we unilaterally inactivated PPC (contralateral to the joystick) by optogenetically stimulating inhibitory neurons in two VGAT-ChR2 mice after they achieved expert performance (Supp. Fig. 5 for behavior quantification in experts, Supp. Fig. 6a-b for map of inactivation spots over PPC). In the first mouse, we saw a drop in accuracy (∼10-15%) where the proportion of joystick movements to the opposite direction increased, consistent with an inability to translate the learned associations into correct movements (Supp. Fig. 6c). In the second mouse, accuracy effects (Supp. Fig. 6d) were modest and individual sessions were variable. However, the reaction times were significantly lengthened (Supp. Fig. 6c-d, inset), suggesting that the mouse compensated by taking additional time to process the stimulus and choose an action.

**Figure 5.**
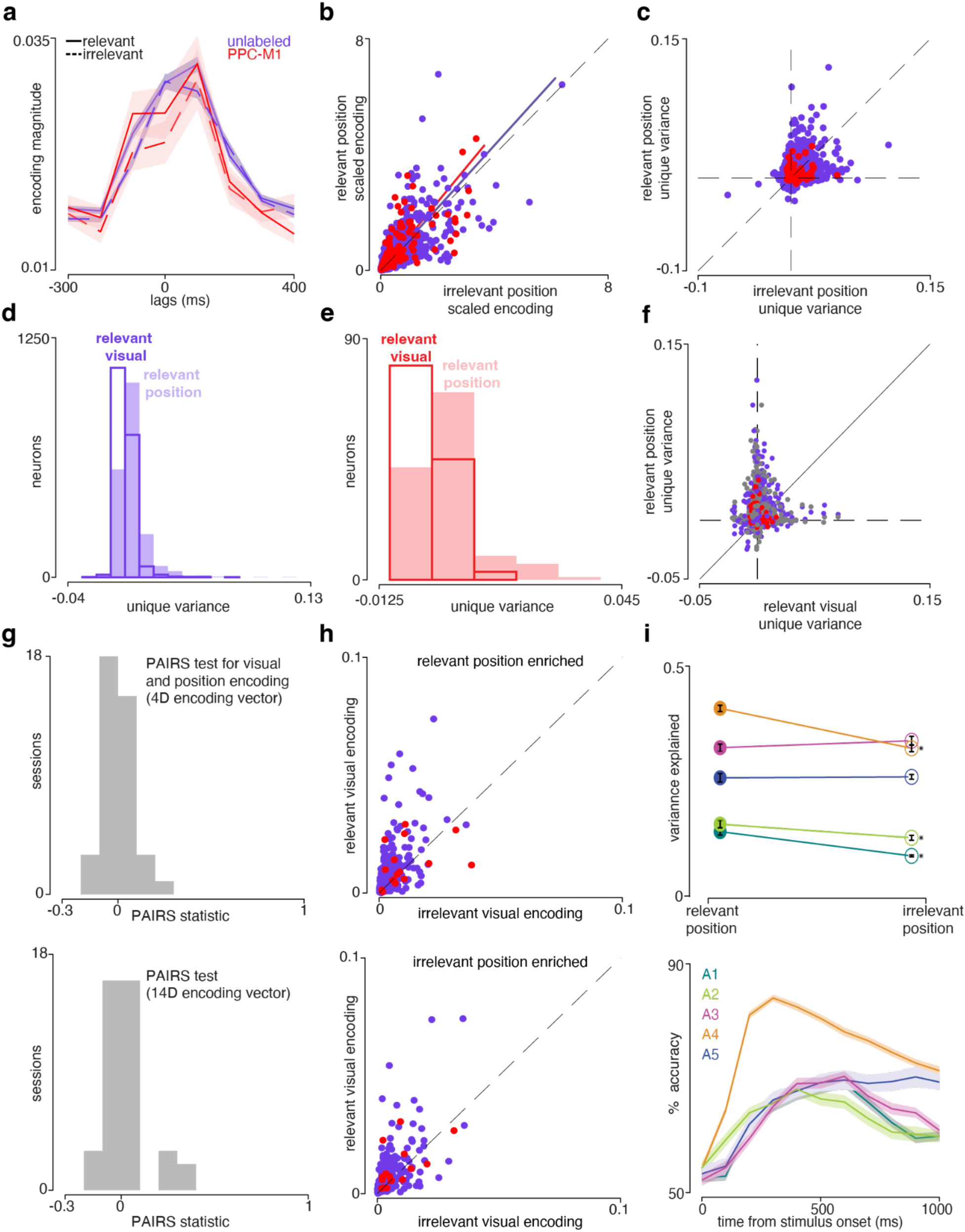
Encoding and decoding of task-relevant and irrelevant movement information in PPC. (a) Population-averaged task-relevant (solid) and irrelevant (dashed) joystick position encoding kernel for unlabeled PPC (purple, n=1,786) and PPC-M1 (red, n=120) neurons. Shaded regions represent the SEM. (b) Single neuron encoding magnitude for the relevant and irrelevant joystick position variables (quadratic mean across lags) for unlabeled PPC and PPC-M1 neurons along with the total least square fit lines. Colors as in (a). (c) Scatter plot comparing the unique variance captured by the task-relevant and irrelevant position variables. Colors as in (a). (d) Histogram of unique variance explained by task-relevant visual (hollow) and task-relevant movement (filled) variables in the encoding model for unlabeled PPC neurons. (e) Same as (d) for PPC-M1 neurons. (f) Scatter plot comparing the unique variance captured by the task-relevant visual and movement variable in unlabeled (purple), PPC-M1 (red) neurons, and a pairing shuffle (gray; x-axis predictors were shuffled). (g) *Top:* PAIRS statistic value for 4D encoding vector (relevant, irrelevant position and visual encoding). The median was not significantly different from zero (p=0.87, sign test). *Bottom:* PAIRS statistic value for 14D encoding vector (all the predictors used in the model). The median was not significantly different from zero (p=1, sign test). (h) *Top:* scatter plot comparing the relevant and irrelevant visual encoding magnitude (unique variance) in relevant movement-enriched neurons (task-relevant movement information > 0.01). *Bottom:* for irrelevant movement-enriched neurons (task-irrelevant movement information > 0.01, bottom panel). (i) *Top:* paired plot comparing the cross-validated variance explained by the instantaneous decoder for task-relevant and irrelevant movement information. Each line is a different mouse. Error bars are across sessions. *Bottom:* decoding accuracy across time for the decoder trained to categorize the direction of task-relevant movement information. Each line is a different mouse. Shaded regions represent the SEM across sessions.

**Figure 6.**
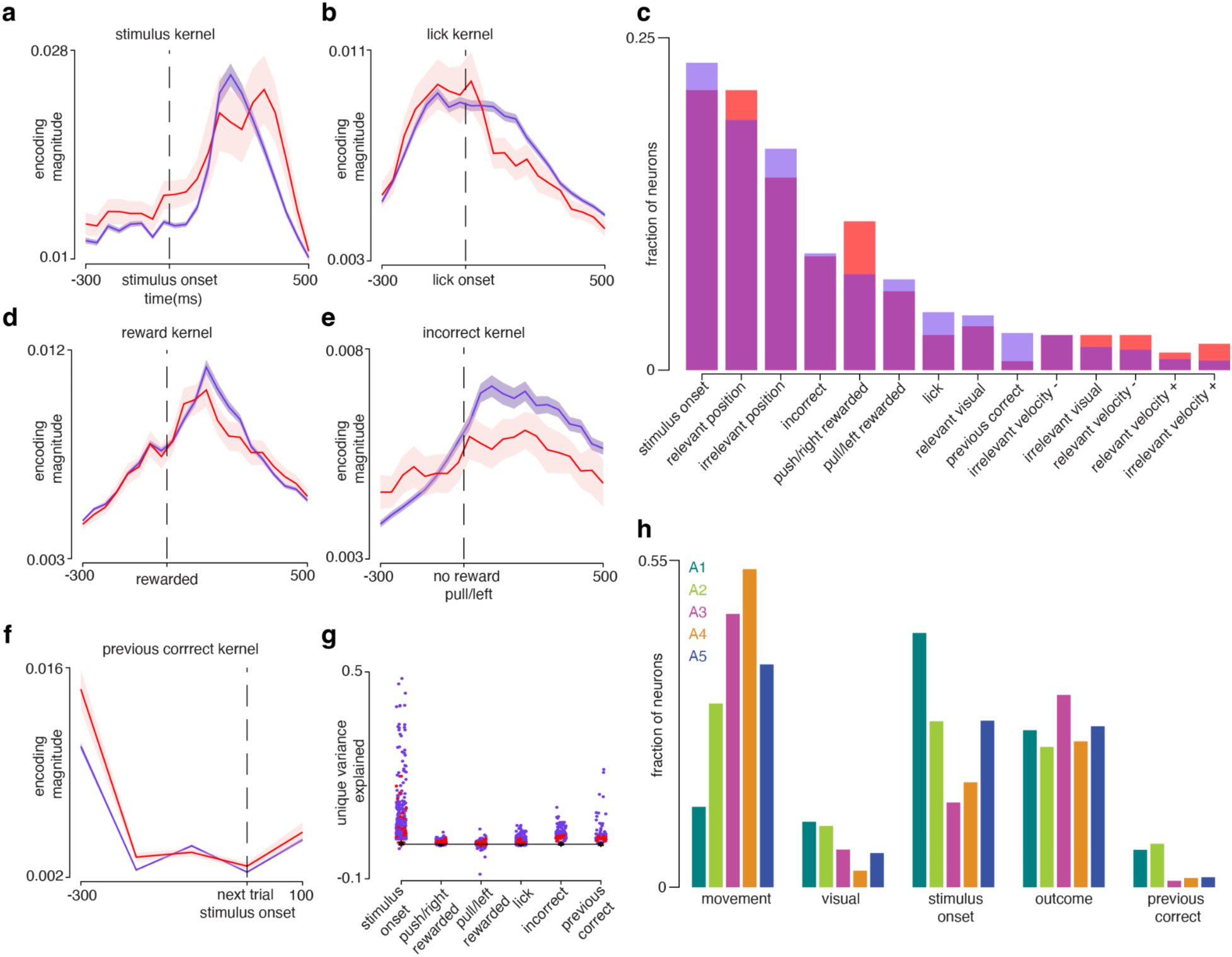
Task-variable encoding in PPC and PPC-M1 during continuous visuomotor control. (a) Population-averaged stimulus onset encoding kernel for unlabeled PPC (purple) and PPC-M1 (red) neurons with the onset time marked by the vertical dashed line. Shaded regions represent the SEM. (b) Same as (a) for the lick onset encoding kernel. (c) Overlaid histograms of the best single predictors for unlabeled (purple) and PPC-M1 (red) neurons. (d) Reward encoding kernel (whether the trial was rewarded). (e) Incorrect encoding kernel (trials where no reward was received). (f) Trial history (whether previous trial was correct) encoding kernel. (g) Distribution of unique variance explained for the event-related model variables for unlabeled (purple) and PPC-M1 (red) neurons. (h) Grouped bar plot of the distribution of best single predictors for each variable type by mouse.

Prior studies have shown that PPC inactivation reduces behavioral biases in addition to impacting visual processing^26,27^. To disentangle the effects of inactivating PPC on sensory vs. “strategy” (superstition) variables, we used a behavioral choice model^36^ to predict the movement from a combination of the stimulus and action history (Supp. Fig. 5e-g for quality of fits). Compared to control trials, models fit to inactivation trials had smaller weights on the stimulus variable for the first mouse (Supp. Fig. 7c), capturing the behavioral deficit. Inactivating PPC had a minimal effect on the movements themselves, with inactivation barely altering the probability of an early joystick release (Supp. Fig. 6c-d).

**Figure 7:**
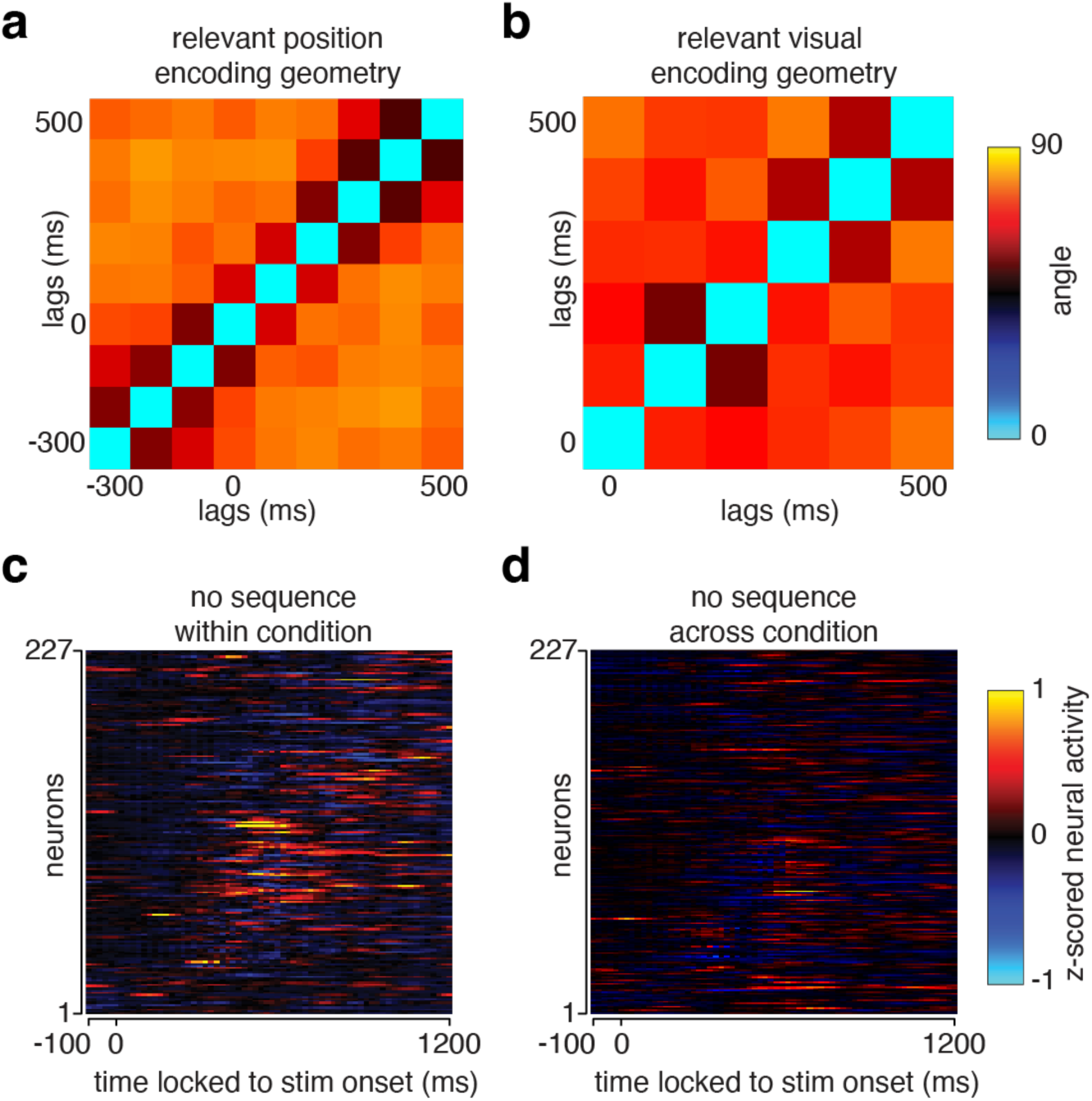
Population encoding geometry in PPC. (a) Angle difference between relevant-visual encoding feature vector at different lags. (b) Same as (a) for relevant joystick position encoding feature vector. (c) Z-scored and trial-averaged residual neural activity (after subtracting activity due to task encoding) from 50% of trials on condition 1 (up-left drift, X-relevant mouse, clean successes), after sorting based on the other 50% of trials. Example session. Note the lack of a diagonal band (sequence) in the plot. (d) Z-scored and trial-averaged residual neural activity from condition 2 after sorting based on condition 1. Same session as (c).

In contrast, inactivation of somatomotor areas produced clear alterations in movement, with mice releasing the joystick early (Supp. Fig. 6e-f). In the first mouse, the errors were entirely independent of trial difficulty, while in the second mouse releases were more common for difficult trials, suggesting a possible interaction (Supp. Fig. 7e,f). Notably, this effect was present after ∼1.5 months of training (Supp. Fig. 1f-g), in contrast with prior findings on simpler joystick tasks where motor cortex dependence ceased with extended training ^37^. Inactivation sessions were randomly interleaved with sessions where the LED was positioned over the headbar, which produced no discernible effects (Supp. Fig. 6g-h, Supp. Fig. 7i-l).

### PPC neurons exhibited heterogeneous tuning, including PPC-M1 neurons

PPC neurons exhibit heterogeneous tuning to a wide range of task-related variables during spatial navigation and multi-sensory decision making^33,8,24,38,34,23^. However, it is unknown what PPC encodes during continuous and closed-loop visual forelimb control; further, it is unknown what signals PPC preferentially represents and/or routes forward in this context. To understand how PPC transforms visual input and propagates it to M1, we imaged the activity of 12,655 neurons in joystick-contralateral PPC neurons in five expert mice (Fig. 3b-e), with labeling to identify whether a neuron sent a projection to M1 (expression strategy shown in Figure 3a; GCaMP6f in most neurons, see Methods for details). Note that unlabeled neurons correspond to a mix of local neurons, neurons that project to targets other than this part of M1, and some number of false negatives due to viral uptake failure or insufficient viral spread. Two-photon imaging data were aligned with the Allen Mouse Brain Common Coordinate Framework version 3 (Allen CCF^39^) using a combination of stereotaxic coordinates and widefield retinotopic mapping (Methods; Supp. Fig. 8). Most of the sampled neurons were in visual area A (A) and we restricted our analysis to neurons that fell within the PPC as defined by refs.^23,30^: higher visual areas AM, RL and area A (Supp. Fig. 9 for animal specific details).

**Figure 8.**
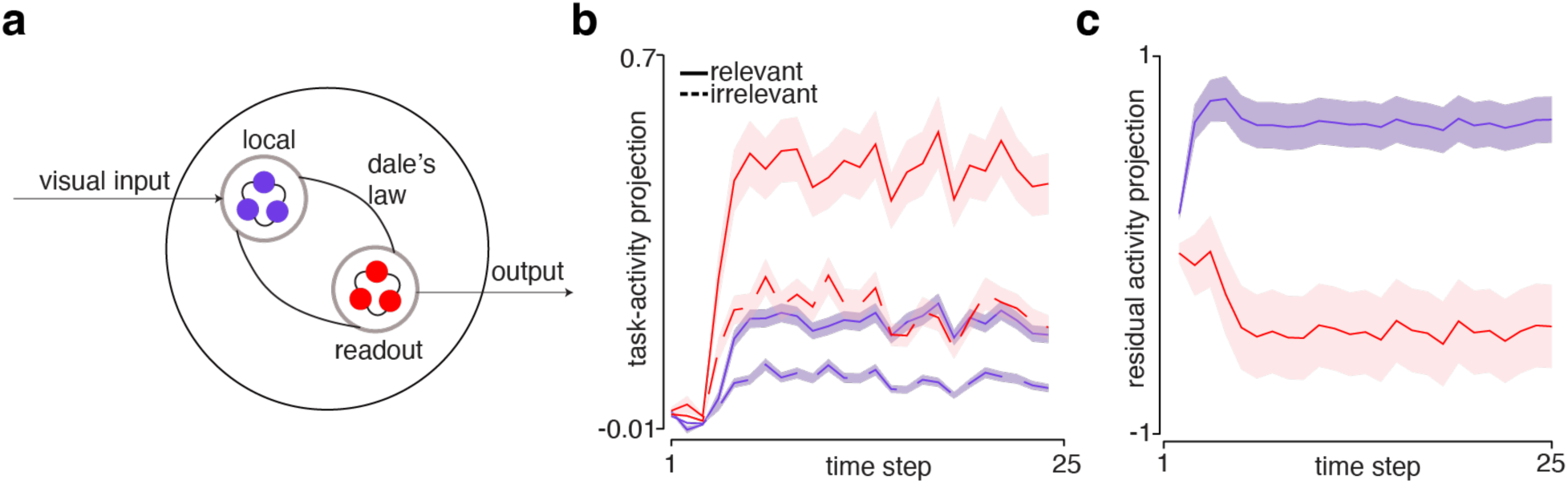
An RNN model trained to act on relevant-visual information. (a) Schematic of the RNN model with segregated local and readout populations emulating the unlabeled and PPC-M1 neurons. (b) Average task-relevant (solid) and irrelevant (dashed) activity projections for local (purple) and readout (red) units in the RNN. (c) Same as (b) for the residual activity.

As a first pass at examining the relationship of neurons’ activity to the task, we trial-averaged each neuron’s response to each of the four quadrants of the visual stimulus (classified using the initial drift direction; correct trials only). Example peri-event time histograms (PETHs) are shown with activity aligned to the stimulus onset and to the final target entry (Fig. 3f-j). Different neurons responded around the times of various behaviorally-relevant events in a trial, such as the onset of the visual stimulus, time of final target entry, or delivery of reward; and presumably-visual neurons could be found with maximal responses to each different drift quadrant. This across-neuron variety in responsiveness to different events and visual properties is consistent with previous reports of mixed selectivity^8,24,40^. The examples shown here highlight some of this variety. One example shows a neuron with a strong but untuned visual response and a weaker but selective response closer to movement (Fig. 3f); another is most active after target entry, perhaps relating to reward (Fig. 3g); a third was active well after reward, suggesting a relationship to behavior in the inter-trial interval; a fourth was selective for both visual drift direction and joystick movement direction (Fig. 3i); and a final example was active before the visual stimulus but stopped firing soon after, perhaps relating to trial history (Fig. 3h)^1,26,27^. At the level of PETHs, PPC-M1 neurons were largely indistinguishable by eye from unlabeled neurons. However, the TextureDrift behavior evoked wide variety in the exact sequence of events on single trials and necessitated a deeper level of analysis.

### Task-relevant visual information is strongly enriched in PPC but not PPC-M1

Three features of the present behavior limit the interpretability of trial-averaged responses. First, single-trial kinematic variability (Fig. 1d-f; Supp Fig. 4g-i), together with the coupling of joystick movements to the visual stimulus, leads to averaging neural activity across trials that featured a wide variety of movements and visual stimuli. Second, this movement-visual coupling poses a challenge in disentangling the effects of visual sensory processing from the effects of movement. Third, the variable duration of the movement epoch leads to the risk of averaging responses during unlike states.

Instead of trying to categorize trials into small and imperfect groupings to permit better averaging, we chose an approach that harnessed the trial-to-trial variability to help distinguish the contributions of different task and behavior variables. Specifically, we fit a linear encoding model to each neuron’s single-trial responses^3–5,38^. Predictors included both continuous-valued variables such as visual drift velocity and joystick position and velocity, and discrete event predictors such as the onset of visual stimulus, onset of the first lick and reward delivery (Methods). To capture any non-instantaneous relationships between neural activity and behavior, we included time-shifted versions of each variable as additional predictors (Methods). Of the 12,655 neurons we imaged in PPC, ∼22% (2,705) met our criteria for being modulated and adequately fit by the encoding model (Methods; Supp. Fig. 10f shows quality of fits).

We considered all the lags of each variable together, forming an “encoding kernel” for that variable. We then examined the encoding kernels in two ways. First, we computed the total magnitude of each kernel, which summarizes how strongly that variable predicted the neuron’s single-trial activity. Second, we examined the shape of the kernel over time, which provides evidence for whether a variable might causally influence neural activity or whether neural activity leads upcoming changes in the predictor. Note that the sign for whether a lag suggests causality depends on whether the neural data or task variables were lagged; here, a negative lag means that neural activity led the task variable, while a positive lag means that the task variable led the neural activity. Crucially, this model permits direct comparison of the encoding kernels for PPC-M1 neurons and unlabeled PPC neurons, to understand what information is propagated directly from PPC to M1.

To examine how visual information is represented in PPC broadly, we asked how strongly the unlabeled PPC neurons encode task-relevant visual information relative to task-irrelevant visual information. Importantly, when we fit the model, we put relevant and irrelevant visual drift components on an equal footing by only including trials with stimuli drifting near 45° in each quadrant (Methods). Population-averaged encoding kernels revealed that task-relevant visual information was more strongly encoded than task-irrelevant visual information in unlabeled PPC neurons (Fig. 4a, purple solid line above dashed; magnitude p=1×10^-32^ by Wilcoxon Rank Sum test; all five animals showed effect individually; Supp. Fig. 10a-e). This analysis treats low-firing-rate neurons as equally important to high-firing-rate neurons, since their ‘importance’ in the circuit is unknown. To see whether there were systematic differences according to firing rate, we scaled the encoding magnitude for each neuron by the range of that neuron’s firing rate as a measure of the possible influence of these encoded variables. This second analysis agreed: task-relevant visual signal was again present at approximately twice the level of task-irrelevant visual signal (Fig. 4c, purple scatter and line).

The encoding kernels of the model provide our best estimate of the importance of each variable to neurons’ firing. The model here was well-conditioned: despite using a task in which the visual stimulus was coupled to the joystick movements, the correlations between predictor variables were modest due to a combination of random initial drift angles, the uncoupled-joystick epoch, and a nonlinearity in the coupling when overshooting the target (Methods). Nevertheless, it is of interest to know how much of the neurons’ firing was uniquely attributable to the visual stimulus. Here, again, task-relevant visual information constituted a larger unique contribution to neural activity than the task-irrelevant visual information did (Fig. 4e; p=6×10^-43^, Wilcoxon Rank Sum test).

What information, then, is transmitted directly to M1? At the level of task-relevant visual information, the existence of highly task-relevant neurons within the unlabeled population suggests that PPC could in principle selectively funnel task-relevant visual information through PPC-M1 neurons to specifically inform movements. Alternatively, PPC-M1 neurons might veridically carry the same mix of signals present in PPC more broadly, performing no further selection. To answer this question, we repeated the same analysis with identified PPC-M1 neurons (red in Fig. 4a,c,e). To our surprise, PPC-M1 neurons carried less task-relevant visual information than the population as a whole (relevant not significantly greater than irrelevant for PPC-M1 neurons; p=0.0062 for unlabeled having a larger relevant-visual vs irrelevant-visual difference than PPC-M1 in kernel magnitude assessed via Wilcoxon Rank Sum test; difference was present for all four animals with labeled PPC-M1 neurons individually; Supp. Fig. 10a-e). This was true as well when using the firing rate-scaled encoding kernels (purple line steeper than red in Fig. 4c), though due to the heavy-tailed distribution of firing rates, did not reach statistical significance (p=0.086, two-tailed neuron identity shuffle; 10,000 shuffles). At least in this pathway and this task, then, PPC underrepresented task-relevant visual information in what it sends directly to M1. These findings are consistent with the linkage principle for visual vs. motor information; but, the loss of task-relevant visual information in the PPC-M1 pathway was not consistent with any hypothesis in the field we know of.

The encoding structure assessed so far captured the independent activity of single neurons. However, neurons act in concert and reformat information at the population level in ways that either limit the overall information or enhance task-relevant information available for readout^41–45,25^. Therefore, despite similar levels of relevant visual information observed in individual unlabeled PPC and PPC-M1 neurons, the coordinated activity across these two populations could represent different levels of information, arguing for a population-level filtering strategy.

To compare the magnitude of relevant and irrelevant visual information decodable at the population level, we trained linear decoders to predict instantaneous visual motion in the relevant and irrelevant direction (see Methods). Consistent with our single neuron encoding results, PPC population activity supported decoding of task-relevant visual information with higher fidelity compared to the irrelevant information (Fig. 4b,d). Next, we built categorical decoders to decode the relevant visual direction (whether the stimulus moved up/down or left/right as applicable) over time. Relevant visual accuracy peaked 250-300ms post stimulus onset (Fig. 4f), reflecting the time course of movement. These population-level analyses support the same conclusion as the single-cell results: task-relevant visual drift information was more strongly encoded in the PPC population than task-irrelevant visual drift information.

### Task-relevant and irrelevant movement information are comparably encoded in PPC and PPC-M1 neurons

PPC neurons encode the upcoming position during spatial navigation^46^, arguing for a ‘higher-level’, more instructional role in movement control. However, whether PPC encodes continuously controlled forelimb movements, and if so at what level it does so, is unknown. Further, in our task, movement variability was predominately in the relevant axis and only this movement was reward related. PPC is more modulated during task performance^20^ and therefore it is possible that PPC would be enriched for task-relevant signals related to movement as well as visual information. Alternatively, PPC could encode both task-relevant and irrelevant movements, consistent with a general state estimate of the animal^34^. This state could then be enriched – or not – for propagation to M1 through PPC-M1 neurons.

From the encoding kernels for joystick position (Fig. 5a), both causal and sensory lags yielded strongly non-zero coefficients, with little difference in the encoding strength between the relevant and irrelevant axes. This was true for both unlabeled (purple lines) and PPC-M1 projection neurons (red lines) and was supported by firing-rate-scaled encoding (Fig. 5b) and unique-variance encoding (Fig. 5c). Results were also similar when using the relevant and irrelevant axes as defined by the task instead of the principal components of movement used here (Supp. Fig. 11a,d).

Both PPC and PPC-M1 neurons encoded relevant movement strongly compared to relevant visual information (Fig. 5d,e). To determine if there was a systematic relationship between encoding of movement and visual motion, we asked if neurons with strong tuning to relevant visual information also tended to exhibit strong tuning to relevant movement. Comparing the unique variance contribution, relevant movement and relevant visual motion were independent from each other (Fig. 5f, gray pairing-shuffle dots do not lie more along the diagonal than purple dots). This was true for the PPC-M1 neurons as well (Fig. 5f, red dots), suggesting they exhibit comparable levels of mixed selectivity. Randomly mixed selectivity was confirmed by a PAIRS test^24^ on the four-dimensional vectors of visual and movement coefficients, which produced only the amount of non-uniformity to the encodings expected by chance (PAIRS statistic values centered at zero; Fig. 5g, top). Similar results were observed when a PAIRS test was run on the fourteen-dimensional encoding vector (including all predictors used to fit the model, Fig. 5g, bottom). To look for any nonlinear effects, we sub-selected the highly task-relevant and the highly task-irrelevant movement neurons (based on their unique variance explained) and examined their visual encoding (Fig. 5h, top and bottom respectively). Again, there was no systematic relationship between the visual and movement encoding.

As above, single-neuron results do not necessarily translate to population-level encoding. We fit population decoders across a range of lags and examined the decoding performance at the optimal lag (see Methods). PPC activity provided reliable decoding of the instantaneous joystick position, both relevant and irrelevant with little or no difference (Fig. 5i, top), consistent with our single neuron results. Next, we built categorical decoders to decode the relevant movement direction (whether the animal moved up/down or left/right as applicable) over time. Relevant movement accuracy peaked around 300-600ms post stimulus onset (Fig. 5i, bottom).

### PPC and PPC-M1 neurons encode numerous task-related variables

To determine how a variety of possible behavioral variables were encoded in PPC during continuous control, we examined the other encoding kernels. First, the stimulus kernel (Fig. 6a) peaked around ∼120ms following the onset of the visual drift, consistent with an external stimulus driving neural responses and providing evidence of minimal cross-talk between visual and movement variables in the model. On the other hand, licking activity (Fig. 6b) peaked before zero, suggesting PPC anticipates or drives the movement. Reward information (both correct and incorrect) was also strongly encoded with a peak following close after outcome onset (Fig. 6d-e). Consistent with prior reports^26,27^, we observed trial history representations that arose shortly after the reward and decayed slowly closer to onset of the next trial (Fig. 6f). To compare the relative strength with which each variable was encoded, we used reduced models (similar to Fig. 4e) to quantify the unique variance contributed by each variable (Fig. 6g). Stimulus onset generally had the strongest contribution, followed by licking and reward, then the task-related visual information, consistent with the conclusion that the visual information encoded in PPC was a small component of the total activity.

In almost all cases, the event-related information was similar in PPC-M1 neurons (red in Fig. 6a-b,d-g). Interestingly, in several kernels, PPC-M1 neurons exhibited a slightly higher magnitude of encoding prior to stimulus onset. This may reflect greater persistent activity from the previous trial. Otherwise, we observed that PPC and PPC-M1 had comparable levels of encoding with very small or no differences in the temporal evolution.

To determine whether PPC-M1 neurons might be more or less mixed in their representations than unlabeled neurons, we examined the distribution of the strongest predictor (Fig. 6c) and observed that PPC-M1 neurons were similar to unlabeled cells (see also Fig. 6h for distribution of the strongest predictor across different categories). This argues that information was structured similarly in these groups, and other than the reduction in task-relevant visual drift information and perhaps the response to errors, PPC-M1 outputs were not specialized to carry any specific subset of signals. Instead, they routed the entire remainder of the local mixed representation with minimal reweighting.

### Population encoding geometry in PPC

In virtual reality navigation tasks, PPC exhibits sequential activity that is thought to encode both the progression along the route and the end-goal^16,33,47^. We therefore asked whether there might be an underlying intrinsic sequence during continuous forelimb control. Alternatively, PPC activity during this task might look very different, consistent with reports that PPC activity reflects task demands^48,49^. To probe the temporal progression of activity, we first examined whether the encoding of neurons changed across time. Specifically, we determined the change in direction of the encoding vectors over different lags (see Methods for details). We observed that both the relevant-visual and relevant-movement encoding vectors changed substantially over just 100ms (Fig. 7a and 7b), reflecting a rapid change in the population of neurons active for a given time period. To determine if these changes constituted an intrinsic sequence, we performed two further analyses. First, we asked whether the order in which neurons activated was conserved across similar trials, by ordering neurons by their activation times on one set of trials and examining the sequence on another set. This was performed after residualizing by the model estimates, to remove ordering due to tuning. However, this neuron order did not generalize to held-out trials from the same condition (example in Fig. 7c), nor across different conditions (example in Fig. 7d). This non-conservation implies that there was not a consistent order by which PPC neurons were recruited across time in this task, beyond the ordering due to tuning for task variables.

### An RNN network model to probe candidate mechanisms for relevant information selection

Thus far, we observed that despite the observed enrichment for relevant visual drift, irrelevant visual information persisted in both the unlabeled and PPC-M1 populations. To better understand whether this preservation is the “natural” solution for a modular circuit or not, we implemented a toy model: a simple Recurrent Neural Network (RNN) with biological constraints trained to solve a simpler version of the task (see Methods; Fig. 8a). Once the network was trained, we identified task-relevant and irrelevant dimensions using categorical decoders trained to decode either the relevant or irrelevant visual direction. In the RNN, task-relevant activity was much more highly enriched in the read-out units than in the local units (Fig. 8b), a marked difference from the recorded neural data. Additionally, the residual activity (all other modulation) was much lower in the output neurons compared to local units (Fig. 8c), inconsistent with the real PPC-M1 and unlabeled neurons having comparable encoding for other variables. Notably, however, some task-irrelevant activity persisted in both the local and read-out population, consistent with the real neural data. This revealed that even a simple network model where irrelevant information did not contribute to the outputs and where no downstream module was available to further filter out irrelevant information, the system did not completely suppress irrelevant information in its outputs. However, the marked differences of the toy model from the real data suggest there may be pressures on the real system to retain at least some components of the task-irrelevant signals.

## Discussion

Despite previous work on the role of mouse parietal cortex in high-level vision and movement planning, little has been known about what signals are propagated to motor areas to flexibly guide behavior. Using a closed-loop visuomotor-control task that strongly engaged PPC in mice, we examined how visual and movement signals are encoded in PPC and propagated to M1. We first characterized the encoding structure in PPC during closed-loop forelimb control. PPC activity was enriched for task-relevant visual signals, which must be learned: X-relevant mice were enriched for X drift, while Y-relevant mice were enriched for Y drift. As in prior work, the encoding of this variable in individual neurons was randomly mixed with other task information such as movement, reward and choice history. Second, we discovered that PPC-M1 projection neurons are similar to other PPC neurons in carrying a broad mixture of signals. Surprisingly, though, these projection neurons carried less of the task-relevant visual activity. Together, these findings suggest that, at least in this task, PPC indeed has a mixed role as both a higher-order visual area and as a high-level movement area, with the movement role predominating in the PPC-M1 pathway.

These findings are consistent with the linkage principle^10,11^, that areas send signals resembling those of their target, for visual vs. movement activity. However, the reason for the reduced task-relevant visual activity in the PPC-M1 pathway is less clear. One possibility is that the relevant-enriched neurons are specific to a different pathway that was not identified here. A number of candidates exist: feedback to other visual areas^7,23,30,31,50–53^ or to pulvinar^54–56;^ neurons that more strongly influence layer 5 activity^7,31^; neurons informing transthalamic projections to motor areas^57^; or direct projections to M2^12,32^. The identity of those neurons therefore remains to be determined. How PPC obtains the enrichment for task-relevant visual signals is also unclear: inputs to PPC might already carry it, such as from pulvinar input that has been linked to visuomotor coupling^58,59^; or this enrichment may be produced locally.

The linkage principle has previously been demonstrated in the early sensory pathways of V1-LM^14^, S1-S2 and S1-M2^15^, and the corticogeniculate pathways in monkeys^60^. Interestingly, however, this result differs from another previous finding, where mouse PPC was found to preferentially send visual signals to M2 and M2 to preferentially send movement signals to PPC, speculatively to avoid “reverberant” activation^13^. Whether the difference with our findings stems from a difference in open-loop vs. closed-loop task, forelimb movement vs. eye movement control, a difference in the PPC pathways to M1 vs. M2, or some other factor is not yet clear.

PPC carrying more movement activity than visual is consistent with a number of prior findings^23,34^ and suggests that mouse PPC may at least sometimes act predominately as a high-level movement area, much like the Parietal Reach Region of PPC in monkey^61–64^. The movement-dominance also echoes results from decision-navigation tasks in mice, where PPC has been found to encode more movement than visual signal^34^.

Notably, the role of a brain area can also vary by task^65–68^. Our task differed from most others used in mice, in that it not only required continuous control of the forelimb in closed-loop with visual feedback but also included an irrelevant dimensions of the stimulus on an equal footing with the relevant one. Further, we did not observe sequential activity often seen in PPC during many decision making tasks. Instead, we observed a rapid change in population geometry, potentially caused by temporally sparse activity but not including an inherent sequence. However, this is consistent with the behavioral outputs that the animal produces. Unlike locomotion which has a reasonable degree of stereotypy, forelimb movements during continuous control are relatively more flexible and variable. Importantly, there are many potential paths towards the same target location. It is possible that any of these aspects of the task further brought out the motor role of PPC and the PPC-M1 pathway.

The implications of mixed selectivity in PPC depend on how readout is performed. Recent evidence combining ensemble holographic stimulation with visual discrimination tasks in mice suggests that readout is performed over the population relatively holistically; manipulating the activity of the most task-relevant-encoding neurons in early visual areas has less impact on behavior than that would be expected from a strongly selective readout strategy^69^. In the RNN that we trained with a greatly simplified analog of the task, task-relevant information was highly enriched in the outputs. This is in stark contrast to our neural results, where task-relevant visual drift information was suppressed in the “outputs” – the PPC-M1 pathway. Presumably, then, in PPC the relevant visual information follows a different route to inform movement, and like task-irrelevant visual information in the model, visual information of all types simply has not been eliminated from the PPC-M1 pathway. This is perhaps surprising given that the mouse brain exhibits much more complete connectivity between areas than the primate brain^29,70,71^. It is possible that the weak but almost complete connectivity structure in mice dictates a less feedforward format to cortical information flow. A recent study in mice showed that both the feedforward and feedback connections between the visual areas V1 and LM carry fine-grained information that serve to build consensus between two areas^72^. Instead of a feedforward chain of refinement steps, then, it may be that the feedforward and feedback signals contribute to joint dynamics that stabilize the required signals in the mouse.

We note two limitations of the present work. First, the genetic strategy used here likely caused calcium indicator to be expressed in inhibitory cells as well as excitatory ones^73^. In contrast, long-range projection neurons are almost all excitatory neurons. Thus, it is likely that the unlabeled population includes contributions from different cell types. If inhibitory cells were for some reason enriched for task-relevant visual signals, this could account for the difference we identified. Relevantly, however, the labeled projection neurons and unlabeled neurons were extremely similar in almost every other way we measured. Second, the visual and movement signals were tightly intertwined in our closed-loop task design. Although a variety of features in our task strongly decorrelated these features, they were causally linked, and it is possible this influenced the nature of the activity in PPC.

Together, these findings support a view of PPC as closely interacting with the motor system, and lay groundwork for further study of signal routing and how function is compartmentalized, bottlenecked and distributed across cortex. Future work to identify the signals routed between other pairs of areas and in other contexts will be essential for determining how brain areas coordinate to use sensory inputs to guide action.

## Methods

### Animal subjects

All animal training and surgery procedures were approved by the University of Chicago Institutional Animal Care and Use Committee. For two-photon imaging experiments, six Ai-148D (TIT2L-GC6f-ICL-tTA2, stock 030328; Jackson Laboratory) transgenic male mice were trained to perform the task (n=4 for the Y-relevant cohort and n=2 for the X-relevant cohort, 8-12 weeks old at the start of training). Five of the six mice were heterozygotes, with the remaining animal homozygous (animal 5/A5). One Y-relevant mouse failed to reach expert performance (>60-70% “clean” success) after 2-3 months of training due to severe bias and was excluded from the study (data not shown). The two-photon imaging data presented here is therefore from five animals (A1-A5). For the inactivation experiments, two male VGAT-ChR2 mice (trained X-relevant) aged 8-12 weeks at the start of training were used. For both imaging and inactivation experiments, mice underwent a single surgery prior to the start of behavioral training. After the surgery, mice were individually housed in a reverse 12-h light/dark cycle, with an ambient temperature of 71.5 °F and a humidity of 58%. Behavioral training sessions and imaging experiments were performed in the afternoon, during the animal’s dark cycle.

### Surgery

Anesthesia, basic surgical preparation and head bar attachment were as described in ref. ^74^. A circular craniotomy was made using a biopsy punch (3, 4 or 5-mm) centered over the left PPC at approximately 2 mm anterior and 1.7 mm lateral of bregma (coordinates from ref. ^16^). In animals with one of the larger craniotomies (>3mm), the window was centered slightly anterior to left PPC to include forelimb M1 (M1-fl). The craniotomy was cleaned with SurgiFoam (Ethicon) soaked in phosphate-buffered solution (PBS). A durotomy was performed with a #15 scalpel and forceps.

Virus (AAV9-CaMKII-Cre, diluted to 1×10^13^ particles per mL in PBS, Addgene stock 105558-AAV9) was then pressure injected (NanoJect III, Drummond Scientific) at 2-7 sites near PPC to induce GCaMP expression. At each location, virus (150 nL) was injected at two depths for sites closer to PPC (250 and 500 μm below the pia) and at one depth (250 μm) for sites near V1 and RL. Each injection took 5 min. The injection pipette was kept in place for another 5 minutes after injection to ensure that viral particles dispersed into the tissue before moving it.

For the labeling of projection neurons from PPC to M1, another virus injection (AAVretro-tdTomato, diluted to 1×10^13^ particles per mL in PBS, Addgene stock 59462-AAVrg) of 150 nL was made in M1-fl at 1.5 mm posterior and 0.4 mm lateral of bregma at two depths (250 and 500 μm from the pia, 1-2 close injection sites). This injection retrogradely labeled the somas of cells that sent projections to M1-fl. In mice 1 and 2, a small craniotomy was made using a dental drill over the left M1-fl to inject the virus. In mice 3-5 M1-fl was accessible within the large craniotomy over PPC.

In mice 1 and 2, the craniotomy was sealed with a 3mm round cover glass glued to a 4 mm round cover glass (Tower Scientific). Prior to the surgery, the two cover glasses were glued to each other using Norbert optical adhesive 81 and cured with UV light. In mice 3-5, the craniotomy was sealed with a single 4 mm #1 round cover glass (Thomas Scientific) glued to a stainless steel cannula (4 mm outer diameter, 3.5 mm inner diameter, 760 µm length, MicroGroup). Prior to the surgery, the cover glass was sterilized using Cidex and glued to the cannula using Zap-A-Gap CA+ slow adhesive (Robart). Windows were then glued in place to the skull first with tissue glue (VetBond, 3M) and then with cyanoacrylate glue (Krazy Glue) mixed with black dental acrylic powder (Ortho Jet; Lang Dental).

For inactivation experiments, two VGAT-ChR2 mice (8-12 weeks) were implanted with a 4 mm #1 round cover glass (Thomas Scientific) centered slightly anterior to PPC to include M1-fl within the window. A single AAVretro-tdTomato virus injection was made in M1-fl as described above.

### Behavioral task

#### Task apparatus

The task was performed by head-fixed, water-restricted mice positioned in a custom 3D-printed translucent acrylic “chair” and controlling a joystick. The chair consisted of two pieces – a seat that supported the animal, and a backrest that positioned their hindpaws and body in a similar posture to that used when grooming. The joystick was positioned under their forepaw with a manual microstage. This position enabled mice to make sufficiently long movements and crucially, groom themselves throughout the session (90-120 minutes). The design was iterated to enable comfortable seating and reliable maneuvering of the joystick and was a crucial element in getting animals to perform hundreds of trials. The left forepaw rested on an eyelet screw paw rest, and both the paw rest and joystick were wired to capacitive touch sensors (Teensy 3.2, PJRC). The joystick itself was a thumb joystick (Apem TS-1D1S00A-1294) with a softer spring installed (Century Spring, part PP-39) and a manipulandum attached: a stainless-steel screw (4/40, 1.5” long) screwed into a 1/2” hex standoff with a jam nut to enable fine length adjustment. To provide grip, a small stainless-steel ball (3/32” diameter) was glued to the top of the screw using epoxy adhesive and coated with nontoxic conductive paint (Bare Conductive water-based electric paint, SKU-0018). The joystick assembly was mounted using a custom 3D-printed acrylic frame. The manipulandum extended ∼2” above the top of the thumb joystick.

#### Trial structure

After mice maintained contact with the paw rest and kept the joystick centered for 300ms, they were presented with a visual texture-like stimulus pattern that drifted in a randomly chosen direction as viewed through a circular aperture. The direction was sampled from a uniform distribution ranging from -60 to +60° from straight in the positive or negative task-relevant direction. The initial drift speed was sampled from a uniform distribution of 41-45°/s. The pattern itself consisted of a black-and-white, camouflage-like image generated similarly to the procedure in ref. ^35^ that was designed to drive mouse visual cortex responses strongly. The only differences from the cited procedure were that the stimulus was spherically distorted here and the pattern was not trimmed. This latter difference meant that the spatial frequencies were not exactly matched to the requested spatial frequencies, but opposite edges were identical and therefore during motion the image could wrap around without discontinuities. A different random texture was generated on every trial. The stimulus was 24 cm in diameter and located 12 cm from the mouse and therefore spanned ∼90° of visual angle. The drift values were an average; a small amount of Gaussian-filtered velocity noise (50ms s.d., with a net standard deviation ∼1% mean amplitude of the average drift) was added to the drift.

For the first 250ms following stimulus onset, the visual drift was decoupled from the joystick. Subsequently, joystick movements were coupled to the visual drift in one dimension (the task-relevant) and mice were required to cancel the relevant drift through a joystick movement of 3.2-6 mm. Movements were more typically 8-12 mm due to some movement in the irrelevant axis and overshooting the required distance. The relevant component of the joystick position added a velocity vector to the visual drift, implicitly specifying a target location in one dimension. The average irrelevant drift remained unchanged throughout the duration of the trial. Mice had to hold the joystick in the target zone for a period of 100ms to obtain a water reward delivered through a blunt needle placed in front of their mouth. Overshooting the target was permitted to ∼15mm, causing the stimulus to reverse its direction in the relevant dimension, with the gain of the joystick-to-visual drift coupling reduced for the overshoot. Overshoot beyond the limit was considered outside the target and the full joystick-stimulus gain was applied. If the animal failed to hold the joystick within the target, they had the opportunity to make corrective movements within the total trial time limit (18 seconds). The inter-trial interval was 3-4 seconds. If the mouse completed a trial successfully, the circular view changed to green, a 4.4Khz sound was played, and the mouse received ∼3.5 µL of water. If the mouse released the paw rest or joystick early or moved in the wrong direction, the circular view changed to blue, a static noise was played, and the trial was aborted without any reward. For some mice, brief incursions into the opposite direction were not punished; for others they were. The task-relevant direction was chosen during the start of training and remained constant throughout the lifetime for a specific animal.

The behavioral task was written in Matlab 2018a using Psychtoolbox 3.0.14. A Teensy 3.2 sampled the external sensors and interfaced with the behavior control software every 17ms. Capacitive touch sensors were wired to the joystick, paw rest and lick spout. The delay between joystick position and visual display change was approximately 37ms. All channels of sensor activity were saved for offline analysis.

#### Decorrelating visual and movement information

To reliably disentangle the visual and movement information in single trials, the task was designed to decorrelate visual drift and joystick movement despite being coupled in the task-relevant dimension. First, the initial drift varied greatly, causing the same joystick position to map to different visual drift on different trials. Second, the nonlinearity in gain when overshooting the target reduced linear correlation.

#### Shaping protocol

After mice recovered from surgery (2-3 days), they were put on water restriction and tapered to 1 mL/day across 3 days. Animals were handled for 5-10 minutes a day and weighed. On the first day, they were head-fixed for 20 minutes to get acclimatized to head fixation and water was provided directly into their mouths through a syringe with a blunt needle spout.

##### Initiation phase

During this phase (1-2 days), mice were first trained to hold the joystick still for a few hundred milliseconds. Once they could do so, we required them to move the joystick by a small distance (3-5mm) in any direction and hold it there for 50ms. They then advanced to having to move the joystick in the correct direction. For this phase, the visual stimulus direction was fixed to drifting straight down, requiring a joystick push for the Y-relevant cohort; or straight left and requiring a rightward joystick movement for the X-relevant cohort. At the end of this phase, the initial fixation period was slowly increased to 300ms.

##### Target magnitude phase

In this phase (3-5 days), mice were trained to make longer movements; they were able to move the joystick to the required target zone and hold it for 100ms. Until the distance solidified, they were only presented with visual stimuli in one direction (drifting straight downward for the Y-relevant cohort, requiring the animals to push the joystick forward, or drifting to the left for the X-relevant cohort requiring the animals to move the joystick to the right). Once mice reliably produced joystick movements for this direction, the opposite direction was introduced with small magnitude. Mice usually took 150-200 trials to explore other movement directions before crystalizing on joystick movements required to cancel the relevant drift in the second direction. For 1-2 days after the introduction of the second direction, the target magnitude was slowly increased until they matched the first direction.

##### Block phase and introduction of the other direction

After mice reliably produced joystick movements in the relevant cardinal directions (push/pull for Y-relevant or left/right for X-relevant), they entered the block phase. To solidify the visual-movement associations and introduce other visual directions, we presented trials in blocks starting with a block size of 150 trials (including error trials) and slowly reduced it over the course of two weeks until the block size was 10 trials. We also slowly increased the range of visual stimuli that were presented in steps of 10° until the target range was reached (±60° in each direction).

##### Random presentation and penalize-opposite phase

This was the longest phase (1.5-2 months), where mice were trained on the final version of the task. On each trial, the direction of the visual stimulus was randomly drawn, and mice were required to produce the relevant cancelling movement within 18 seconds. For the first two weeks after randomization, they were free to move in the opposite direction and make corrective movements to get a reward. During this period, we slowly reduced the acceptance window of opposite direction movements. At the end of two weeks, animals were penalized for moving in the wrong direction first with a time-out of 5s. Mice continued to train until they reached expert performance and performance level saturated (>60% correct direction in the first attempt, “clean” success). After performance plateaued, the penalty time was reduced to 1.5s. Imaging data presented here were collected ∼2.5-3 months after the start of behavioral training (Supp. Fig. 1).

### Optogenetics

For the optogenetics experiments, two male VGAT-ChR2 mice (stock 014548; Jackson Laboratory) were trained with X as the task-relevant direction. After mice achieved expert performance (Supp. Fig. 1f-g; Supp. Fig. 5), inactivation experiments were performed over the course of 3-4 weeks using the epifluorescence path of the two-photon microscope. On each session, light path location was chosen to be centered near PPC, or the M1-fl / S1 border. Area centers were identified through a combination of retinotopic mapping, projection neuron expression pattern, and stereotaxic coordinates (Supp. Fig. 6). The stimulation protocol consisted of delivering a single 3000ms light pulse (17 mW power with a step onset and 100ms linear ramp down). The spot of light was approximately 1.3 mm in diameter at the focal plane. Stimulation was delivered beginning 300ms before the onset of the visual stimulus. The stimulation probability was fixed at 20% of trials for all inactivation sessions. A machined aluminum cone was fitted over the end of the objective and sealed to the head bar with a ring of gray butyl rubber headlight sealant to prevent leak of the visible blue light. Inactivation sessions were interleaved with headbar control sessions where the light path was positioned over the acrylic covering the head implant.

### Retinotopic mapping using widefield imaging

The widefield macroscope setup was as described in ref. ^74^. Imaging was performed at 30 frames/s with blue illumination focused 0-500 µm below the brain surface. For retinotopic imaging experiments, mice were headfixed in a rig different from the behavioral rig for contextual separation. Retinotopic mapping protocol was similar to procedures described in ref. ^3^, with moving, checkerboard bars swept at 0.5 Hz displayed across one monitor in front of the animal and one to the contralateral side, both oriented vertically and positioned ∼19 cm from the eye. A custom 3D-printed black plastic cone, sealed to the implant with gray butyl rubber headlight sealant, was used to block the light from the monitor entering the window. Processing of the data was similar to ref. ^3^, with some adjustment of smoothing parameters for each animal to maximize map clarity.

### Behavior quantification

#### Trial screening

The following trial types were removed from each session prior to use in any analysis: breaks in fixation, releasing the forepaw from the joystick for more than 300ms during the course of the trial, long response times (>3000ms) and the first two trials. Overall, this removed 2-10% trials in a session.

#### Psychometric functions

A trial was considered a “clean success” if the animal did not move in the wrong direction by the distance required to succeed in the correct direction at any point before the end of the trial. Confidence bounds for each group were computed using the Wilson binomial proportion confidence intervals.

#### Position histogram

To quantify the variability of movement within a session (Fig. 2a,d), joystick data for all trials in a window of -50 to +200ms from target entry were interpolated to 20ms and smoothed with a Gaussian (10ms s.d). The smoothed joystick position was then binned prior to computing the contours.

#### Reaction time

Since the target distance varied across trials, in order to make reaction time (RT) measurements comparable across targets, we took the smallest target distance in a session as a reference threshold. RT was taken as the time the animal crossed this threshold with the joystick for the first time on the trial.

#### Choice model

We used the probabilistic choice model developed by ref. ^36^ to quantify the effect of sensory and non-sensory variables on movement choices. Joystick movements were binarized based on the direction of the first movement, which was taken as the first time the joystick position left the central fixation circle. For optogenetics experiments, for each mouse we fit one model to all control trials across all sessions, and a second model to all inactivation trials across all sessions (Supp. Fig. 5e-g, model fit quality; Supp. Fig. 7, model weights). For imaging experiments, only a single model was fit for each mouse (Supp. Fig. 4a-f). In each trial, the binarized decision *z*(*t*) was modeled as depending on the weighted sum of the signed relevant visual stimulus *s*(*t*) on the same trial, two “strategy” terms (success and failure on the previous trial, *s*(*t*-1) and *f*(*t*-1) respectively) and an overall bias term *b*_0_. The probability ratio of choosing one condition vs. the other was then given by

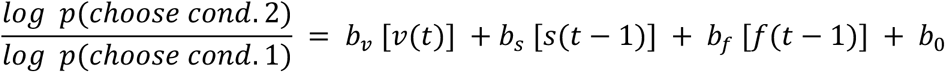

where *v*(*t*) was positive for condition 1 (push/right for Y-relevant/X-relevant) and negative for condition 2 (pull/left). The models were fit with logistic regression.

### Two-photon calcium imaging

Calcium imaging procedures were nearly identical to refs. ^74,75^. Briefly, imaging was performed using a Neurolabware 2p microscope running Scanbox 4.1 and a pulsed Ti:sapphire laser (Vision II, Coherent). Depth stability, ROI curation, deconvolution and time-locking procedures were identical to ref. ^74^. The sole difference was that a machined cone enclosed the end of the objective and was sealed to the implant to block the light from the visual stimulus display as described in the Optogenetics section above.

#### Projection neuron identification

Projection neurons were manually identified using the mean red channel image acquired during 2p imaging. The corresponding ROIs from the green channel were then identified.

#### Peri-event time histograms

PETHs were computed aligned to three different events: onset of the visual stimulus, time of last target entry and reward delivery. Trials were divided into four quadrants based on the direction of the drifting visual stimulus. Neural data for all trials were interpolated to 10ms and smoothed with a Gaussian (35ms s.d). Stimulus-locked PETHs included a window of -200 to 300ms from stimulus onset, and target entry-locked PETHs included a window of -200 to 300ms relative to the event.

#### Aligning FOVs onto the Allen CCF

Alignment procedures for the FOVs were identical to ref. ^74^. Briefly, each 2p FOV was aligned based on the vasculature to an image of the entire window captured via widefield imaging. In imaging mice, phase maps from retinotopic mapping were used to identify V1 and the higher order visual areas AM and RL, and to therefore align the imaged window to the Allen CCF (Supp. Fig. 8-9). Epifluorescence images of the entire window were acquired (MVX10, Olympus) to identify the locations of tdTomato expression for 6/7 mice. Along with injection landmarks, the red “hotspots” served as additional references for alignment (not shown) in the imaging cohort, and for identifying focal spots for activation and alignment in the optogenetics cohort. Two-photon images from the cortical surface from each session were then scaled to their measured sizes and aligned to the registered vasculature via rigid transformations. This series of transformations produced global coordinates for each ROI from its location within the 2p FOV.

### Linear encoding model

#### Model construction

The encoding model was similar to that in ref. ^3^. The design matrix was constructed from a number of binary event-related variables and several analog task-variables. In both cases, the variable was captured by a predictor vector and time-lagged copies of that vector, in order to capture the relationship at multiple temporal offsets. The event-related variables included predictors for the onset of the visual stimulus, first lick, reward and trial history. Reward was encoded with a set of three predictors: condition 1 correct, which was 1 when the trial belonged to condition 1 and was rewarded, 0 otherwise; condition 2 correct, which was 1 when the trial belonged to condition 2 and was rewarded, 0 otherwise; and incorrect, which was 1 when the trial was unrewarded and 0 otherwise. For stimulus, lick and reward, the event-related predictor consisted of a binary vector containing a pulse (a value of one) at the time of the event. The trial history (or “previous correct”) variable consisted of a step function (1 for the duration of the inter-trial interval and beginning of trial until stimulus onset when the previous trial was rewarded and 0 otherwise; 0 thereafter). The time-shifted copies included both forward and backward lags for all of these event variables. For stimulus, reward and lick variables, the lags were spaced at 40ms (spanning -300ms to 500 ms, locked to stimulus, outcome or first lick respectively). For trial history, the vectors were spaced at 100ms (spanning -300 to 100ms, locked to stimulus onset). The wider spacing for trial history was chosen because of the longer autocorrelation of the variables.

For analog variables, 2D joystick position, 2D joystick velocity, and 2D visual drift were included. Presumably due to biomechanics and the lack of pressure to avoid the irrelevant axis, the largest axis of joystick movement did not align perfectly with the nominally relevant axis. To quantify the two axes of joystick movement, we therefore used the two principal components of the movements instead of the external X and Y axes. Results were similar if we used external X and Y instead (Supp. Fig. 11a,d). Because joystick velocity explained little variance, in an effort to enable fitting asymmetries in how the brain encoded different directions, for this variable only we fit separate positive and negative kernels. In practice this made virtually no difference to fits. Since the analog variables had long autocorrelations, they were spaced at 100ms. Joystick data and visual drift were linearly resampled to 20ms bins. Joystick data was smoothed with a Gaussian (10ms s.d). Joystick velocity was computed by numerical differentiation of joystick position followed by smoothing with a Gaussian (10ms s.d.). The time-shifted copies included only forward lags for visual drift and both forward and backward lags for other analog variables (joystick position and velocity).

To make best use of the data, neural data in the entire trial were included, starting from the 200 ms of initial hold time required to start saving behavioral data (typically ∼300ms before the onset of the stimulus) to 1500ms following trial outcome. Neural data were resampled into 20ms bins and smoothed with a Gaussian (10ms s.d.), then Z-scored.

All columns of the design matrix for each variable were mean subtracted, then normalized by the standard deviation of the unlagged predictor column for that variable. The model was fitted using Ridge regression. The regularization hyperparameter was chosen using Marginal Maximum Likelihood estimation refs. ^3,76^.

#### Trial screening

For all analyses quantifying task-relevant and irrelevant encoding, but not for other analyses of the encoding model, only “diagonal” trials were used – trials where the drift angle was 45 ± 15° in any quadrant. This ensured that for these potentially sensitive comparisons, equal amounts of initial drift were present for the relevant and irrelevant axes. This resulted in discarding approximately half the trials for these comparisons. Results were similar in all cases when analyses were repeated without this trial selection (Supp. Fig. 11b,e). For analyses where all angles could be used, 363 trials were included per session on average. For analyses restricted to diagonal trials, 187 trials were included per session on average.

#### Neuron selection

As a simple criterion for including a neuron for analysis, we required a minimum number of “events” (non-zero deconvolved values). To be included in fitting the encoding model, a neuron had to have at least 100 total events across the session, considering only the portion of each trial starting 200ms prior to stimulus onset and ending 1000ms after reward delivery.

#### Checking design matrix conditioning

To ensure the model was well-conditioned and would produce stable fits, we examined the cumulative subspace angles ref. ^3^. This method finds the angle of each column of the matrix to the subspace spanned by all previous columns of the matrix. Across all sessions, the most collinear variable was 22° from the subspace of previous columns. The average value was 67°.

#### Variance analysis

Total explained variance for the full model was obtained using ten-fold cross-validation. Neurons whose cross-validated fraction variance explained was greater than 0.01 were included for further analysis. To compute the unique explained variance by individual model variables, reduced models were fit where all variables except the specified one were used. The difference in explained variance between the full and the reduced model yielded the unique contribution of that model variable, ΔVE as described in ref. ^3^. The best predictor for each neuron (Fig. 6c,h) was the model variable with the largest unique explained variance for that neuron.

#### Encoding kernels

For each neuron, the encoding kernel for a given variable was the vector of model coefficients for all the predictors corresponding to that variable – that is, across all the fitted lags. Each model coefficient was obtained by averaging the weights over cross-validation folds. To assess whether the encoding kernels were significantly different from noise, the model was refit 10 times to the time-shuffled activity of each neuron. Neurons were deemed to have significant kernels if the real kernels were ≥3 s.d. of the distribution of the shuffle kernels (see Supp. Fig. 11c,f for comparisons of relevant visual and position kernels with estimates from shuffle models). This test was performed separately for each model variable, leading to differences in the number of neurons contributing to each population kernel. Average population encoding kernel magnitudes were estimated by computing the mean across the absolute values of the individual kernels.

#### Encoding magnitude

To compute a summary value of the encoding magnitude for each model variable, we computed the quadratic mean (root-mean squared) across all the lags for each neuron. Computing the sum of the absolute values of the coefficients across all lags or the maximum absolute encoding value yielded similar results. For each neuron, the encoding magnitude reported in Fig. 4c (visual) and Fig. 5b (movement) was scaled to reverse the effect of Z-scoring the firing rate, by multiplying by the standard deviation of the vector of smoothed single-trial firing rates.

#### Encoding geometry

To determine how population activity patterns changed over time (Fig. 7a-b), we compared the population encodings of a variable at different time lags. To do so, for each time lag, a neurons × 1 vector of the encoding coefficients was made per session. The angles between these vectors were then computed. To obtain a mean estimate, the angles were averaged across all sessions.

### Time-varying decoder analyses

To quantify the total amount of relevant and irrelevant information available at the population level, we built instantaneous decoding models using Ridge regression methods described in ref. ^3^ and categorical decoding models using linear Support Vector Machines (SVMs). Trial selection procedures were identical to the encoding model. Cross-validation, partitioning and bagging weight procedures were similar to ref.^74^.

To decode instantaneous visual drift, we used the period where visual drift and joystick movement were uncoupled (0 to 250ms after stimulus onset). In this period, task-relevant and irrelevant visual drift had similar magnitude. The criteria to include neurons for use in the decoding models was similar to those used in the encoding models, with the exception that neurons were chosen based on their activity levels in the period 0 to 1000ms from stimulus onset, and 300 total events were required. To capture the temporal progression of the relationship between neural activity and behavior, fitting was performed separately at each lag using a sliding window (neural data in a window of 200ms, slid 20ms for each time-step, spanning 0 to 1000ms). The cross-validated total variance explained was computed at each lag and the best lag was chosen based on the maximum value of the total variance explained. To decode instantaneous joystick movement, neural activity from stimulus onset to reward delivery was included.

#### Categorical decoding

For visual category decoding (Fig. 4f), the direction of the relevant drift (up/down for Y-relevant; left/right for X-relevant) was decoded from neural activity (0 to 1000ms following stimulus onset) using logistic regression on sliding windows averaged over 100ms. For movement category decoding (Fig. 5i, bottom), binarized joystick movement labels (similar to the choice model) were decoded. Success and failure trials, and push/pull (or left/right) trials, were balanced across all training folds.

### Controls to detect sequential activity during trial progression

To test for sequential activity (Fig. 7c-d), we first computed the residual activity for each neuron on each trial by subtracting the model estimate. This enabled determining if the change in geometry was caused by a sequenced component that was not captured by the model. We next sorted the trial-averaged residual activity for one condition (e.g., leftward trials for a X-relevant mouse) using half of the data, then applied the same sorting to the held-out trials for that condition. The sorting did not generalize to the held-out trials from the same condition (Fig. 7c). We also tested whether the sequence generalized across conditions. To do this, we computed the neuron order using trial-averaged activity from one condition and applied this sorting to the other condition. Again, the sorting order did not generalize (Fig. 7d) suggesting that there was no consistent structure to the order in which neurons were activated across time during continuous forelimb control.

### Task-trained RNNs

A toy recurrent neural network model was trained to solve a simple version of the task. The network received a single input (angle) and was trained to reproduce a negative value of the relevant direction magnitude. The network was trained using the pyTorch and Lightning packages using the Adam optimizer. Since long-range connections in cortex are almost all excitatory, the network was constructed with some constraints. Inputs were restricted to a subpopulation that contributed zero weights to the readout, and all input weights were positive. A separate readout population contributed strictly positive weights to the readout, but note that the outputs could still become negative because of bias contributions. Dale’s law was maintained in the recurrent weight matrix updates. The network was trained until it reached an accuracy of ∼70%. Decoding dimensions were found for both relevant and irrelevant signals using logistic regression on the binarized categories. Single-trial activity was projected onto these decoder dimensions to compare the activity between the local and readout neurons. Residual activity was determined as the single trial activity that remained after subtracting the task-related activity.

## Acknowledgements

The authors thank S. Salimian for animal care assistance, C. Ellithorpe for mouse illustrations, D. Sabatini and M. Greaney for work on the behavioral setup and S. Musall for widefield imaging code. This work was funded by NIH-NINDS R01 NS121535 (MK), the Simons Foundation grant 876393SPI (MK), the Whitehall Foundation grant 2019-12-111 (MK), the NSF-Simons National Institute for Theory and Mathematics in Biology via grants NSF DMS-2235451 and Simons Foundation MP-TMPS-00005320 (MK), the Sloan foundation (MK), the Pritzker fellowship for Neurosciences (PR), NIH T32 NS121763 (HG), and The University of Chicago.

## Author contributions

P.R. performed the surgeries, trained the mice, acquired the two-photon imaging and optogenetics data with assistance from H.G., and implemented the analyses. P.R. and M.K. designed the analyses. H.G. aligned the data to the atlas. M.K. conceptualized the project, supervised all aspects of the project, designed the experiments and behavior, built the setups with assistance from H.G. and P.R., built and performed the widefield imaging with assistance from P.R., and designed the data preprocessing. P.R. and M.K. drafted and edited the paper. M.K. provided funding.

## Supplementary figures

**Supplementary figure 1.**
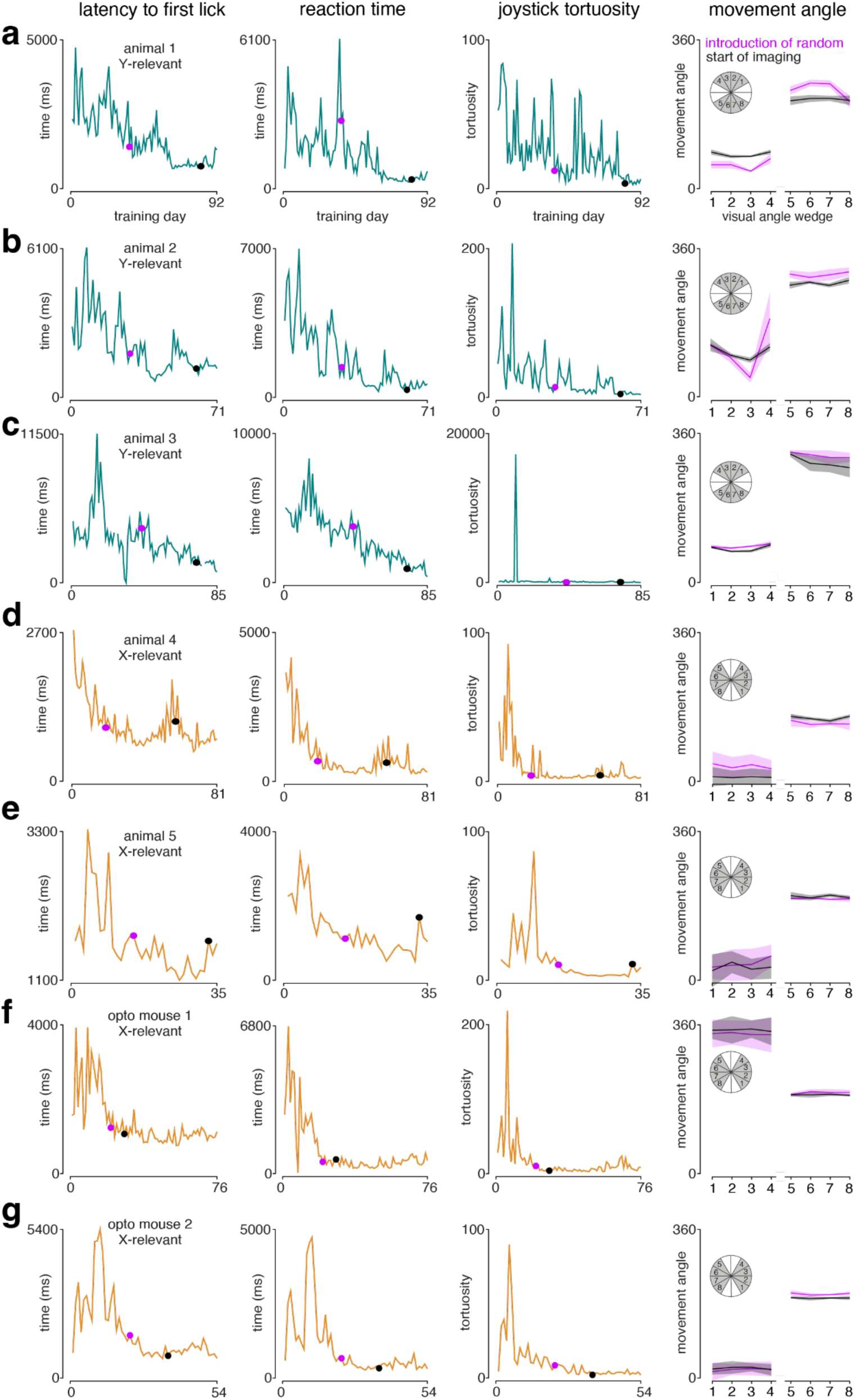
Learning across time. (a) First panel: latency to first lick across training sessions for mouse 1. Magenta dot indicates the day when visual motion direction presentations were made random. Black dot indicates the day when imaging was started. Second panel: reaction time across training sessions for animal 1. Third panel: line plot of joystick tortuosity across training sessions. Fourth panel: joystick movement angle as a function of visual direction on the first day of random presentations (magenta) and first day of imaging (black) for animal 1. Inset: visual direction grouping schematic. Gray indicates the wedges from which visual directions were sampled. White denotes directions that were not sampled in the task. (b-e) same as (a) for animals 2-5, respectively. (f-g) same as (a) for animals from the optogenetic inactivation cohort. Black dots for the inactivation cohort denote the first day of inactivation experiments.

**Supplementary figure 2.**
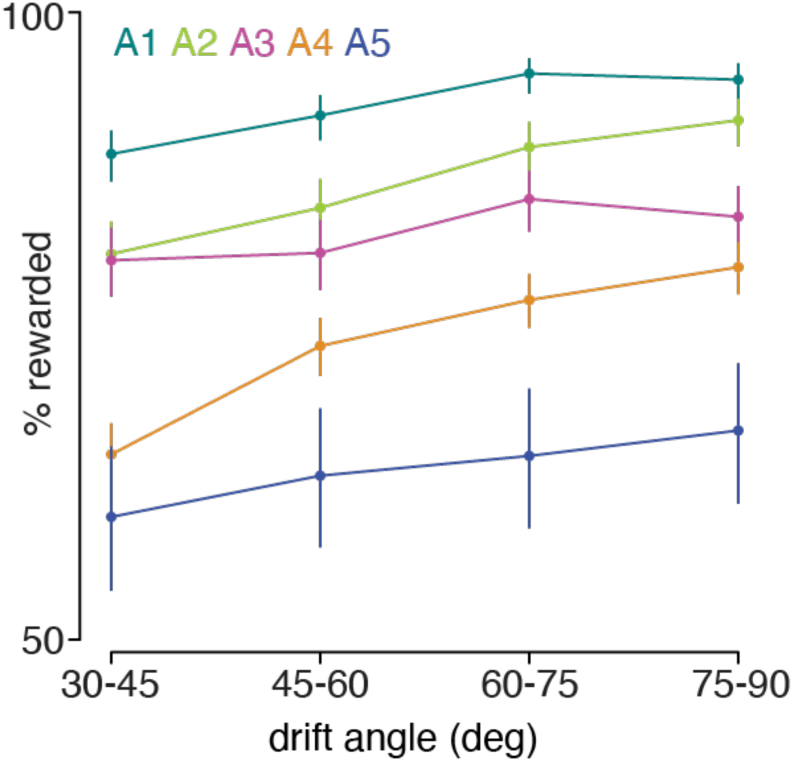
Percentage of rewarded trials as a function of visual drift angle.

**Supplementary figure 3.**
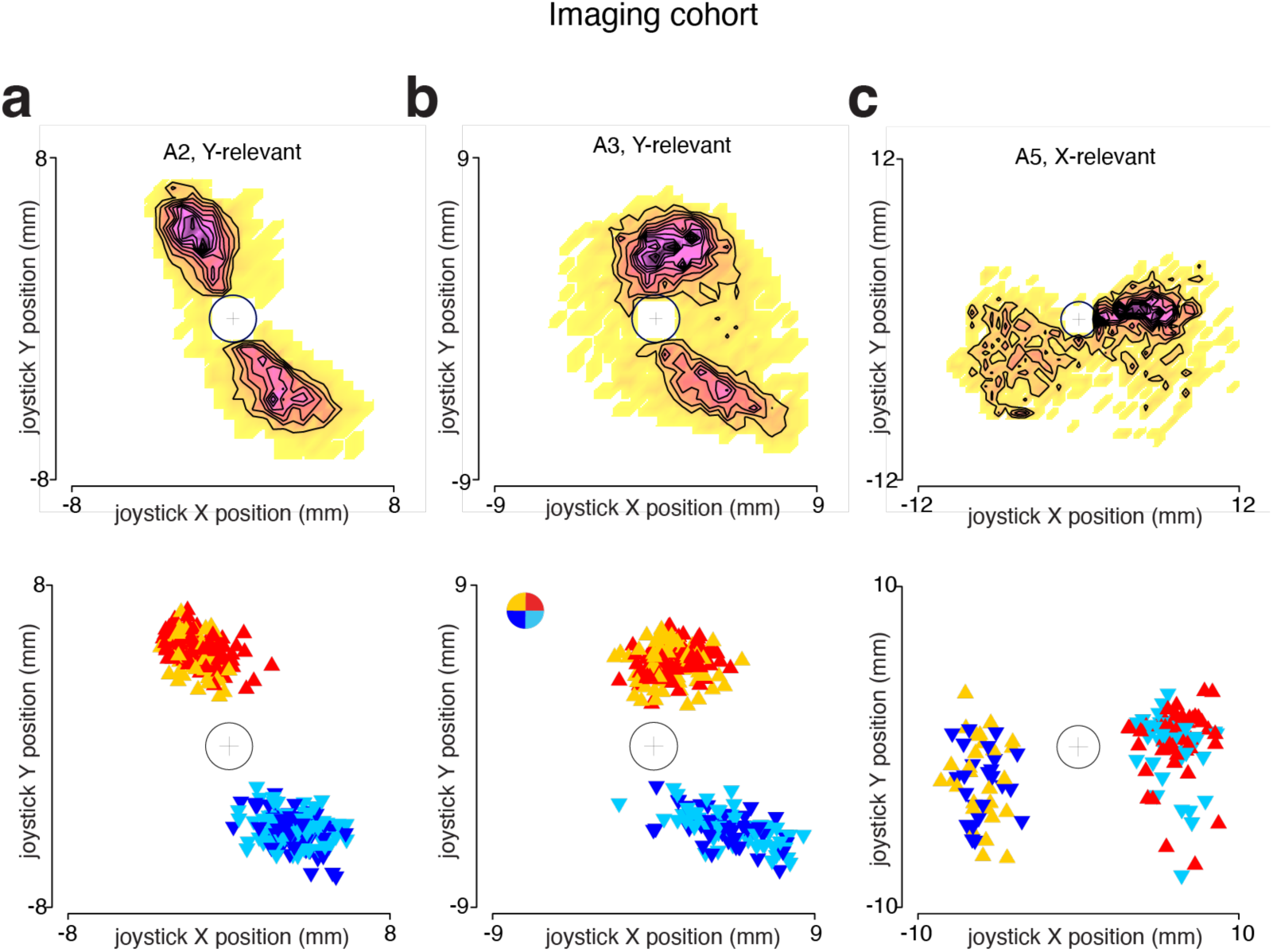
Joystick position during expert behavior. (a-c) Top panels: Binned joystick position (with contours) in a session from mouse A2, A3, A5; central fixation zone (white circle) excluded. Bottom panels: joystick position at the outcome sample for different trials, colored based on the quadrant of the visual drift. Inset color wheel shows the color key by quadrant opposite to the drift direction.

**Supplementary figure 4.**
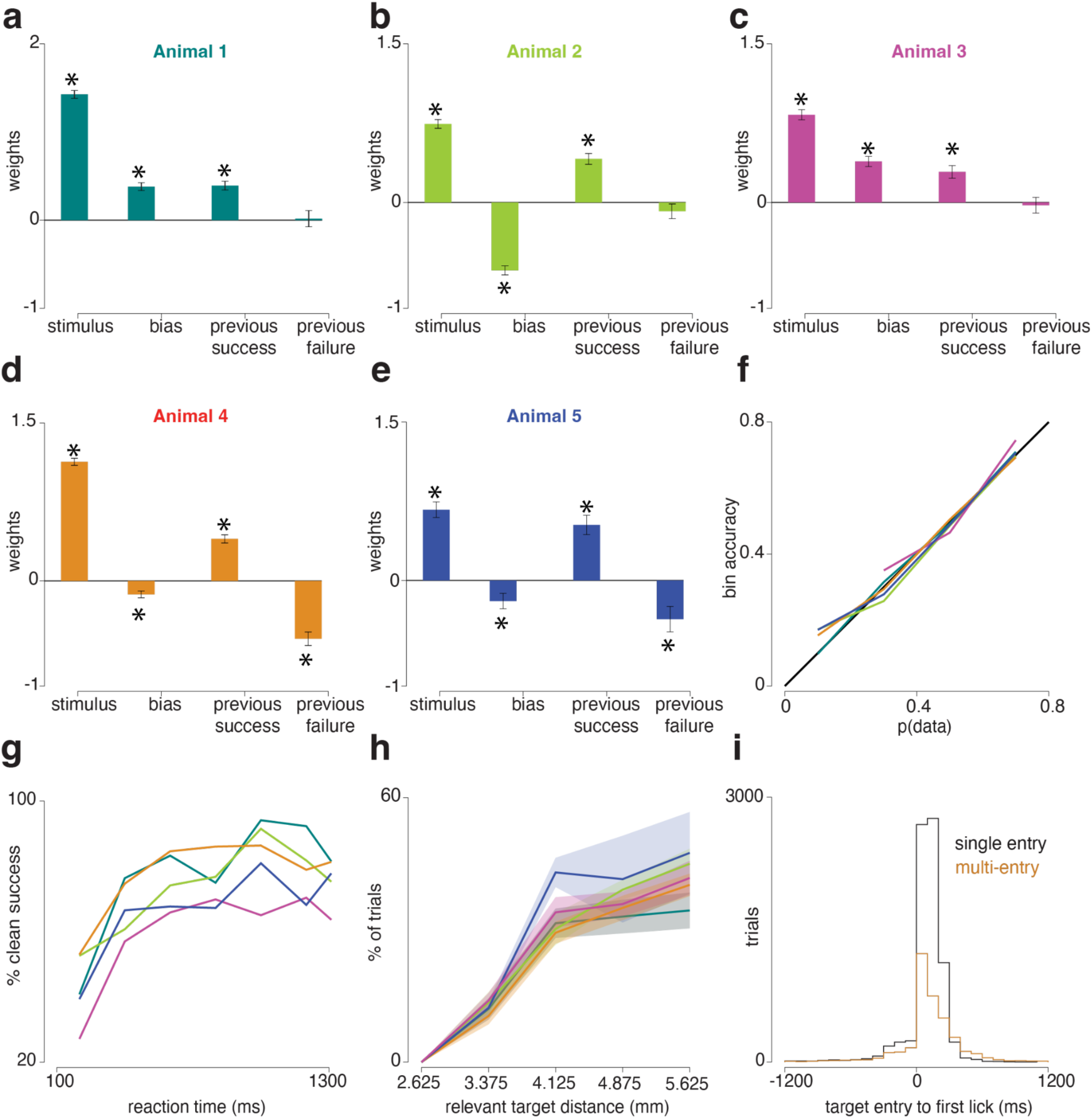
Choice models for the imaging mice and quantifications of behavior. (a)-(e) Estimated weights from logistic regression choice model for the stimulus (signed relevant visual information) and strategy variables (bias, previous success and previous failure) aggregated for each mouse. Trials were concatenated across sessions to fit a global model for each mouse. SEMs were obtained from the model fit. Stars indicate p<0.05, obtained from the model fit. (f) Probability of data vs. estimates from the model. Nearly-straight lines imply minimal nonlinearity unaccounted-for. (g) Proportion of clean successful trials as a function of reaction time. Note that the restriction to clean movements results in a success rate below chance for short RTs, because early movements were less likely to be clean and only about half of trials were successful due to guessing. (h) Proportion of trials with multiple entries into the target zone as a function of target distance. (i) Histogram of time difference between the latency of first lick and target entry.

**Supplementary figure 5.**
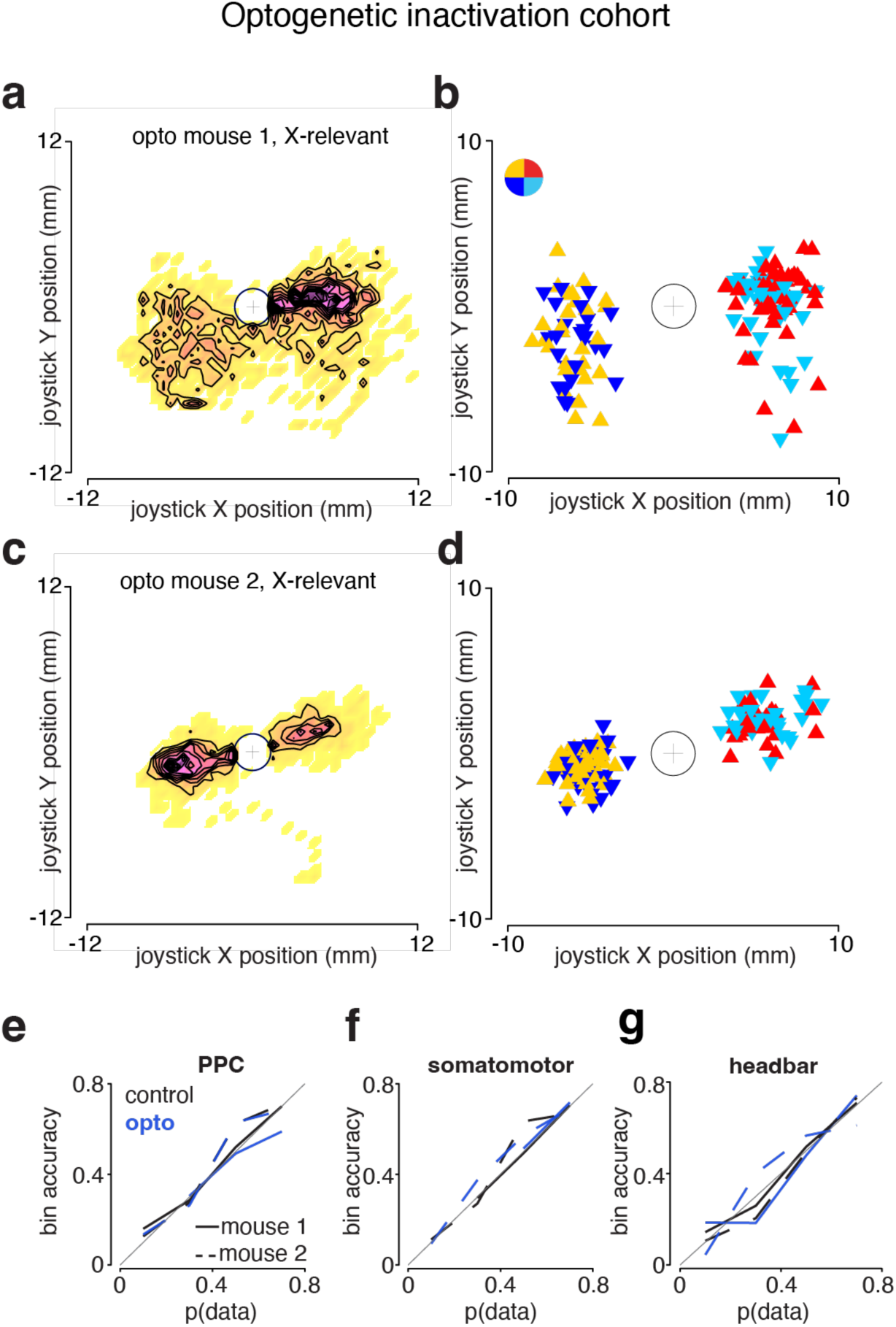
Joystick position during expert behavior and choice model fit quality for optogenetic inactivation cohort. (a) Binned joystick position (with contours) in an example session (prior to the start of inactivation experiments) from optogenetic inactivation mouse 1 (opto mouse 1); central fixation zone (white circle) excluded. (b) Joystick position at the outcome sample for different trials, colored based on the quadrant of the visual drift in opto mouse 1. Inset color wheel shows the color key by quadrant opposite to the drift direction. (c) Same as (a) for optogenetic inactivation mouse 2 (opto mouse 2). (d) Same as (b) for opto mouse 2. (e) Probability of data vs. estimates from the choice model fit to control (black) and inactivation (blue) sessions over PPC for opto mouse 1 (solid) and opto mouse 2 (dashed). Nearly-straight lines imply minimal nonlinearity unaccounted-for. (f) Same as (e) for inactivation sessions over somatomotor areas (S1-fl, S1-hl, M1-fl). (g) Same as (e) for control sessions over the headbar.

**Supplementary figure 6.**
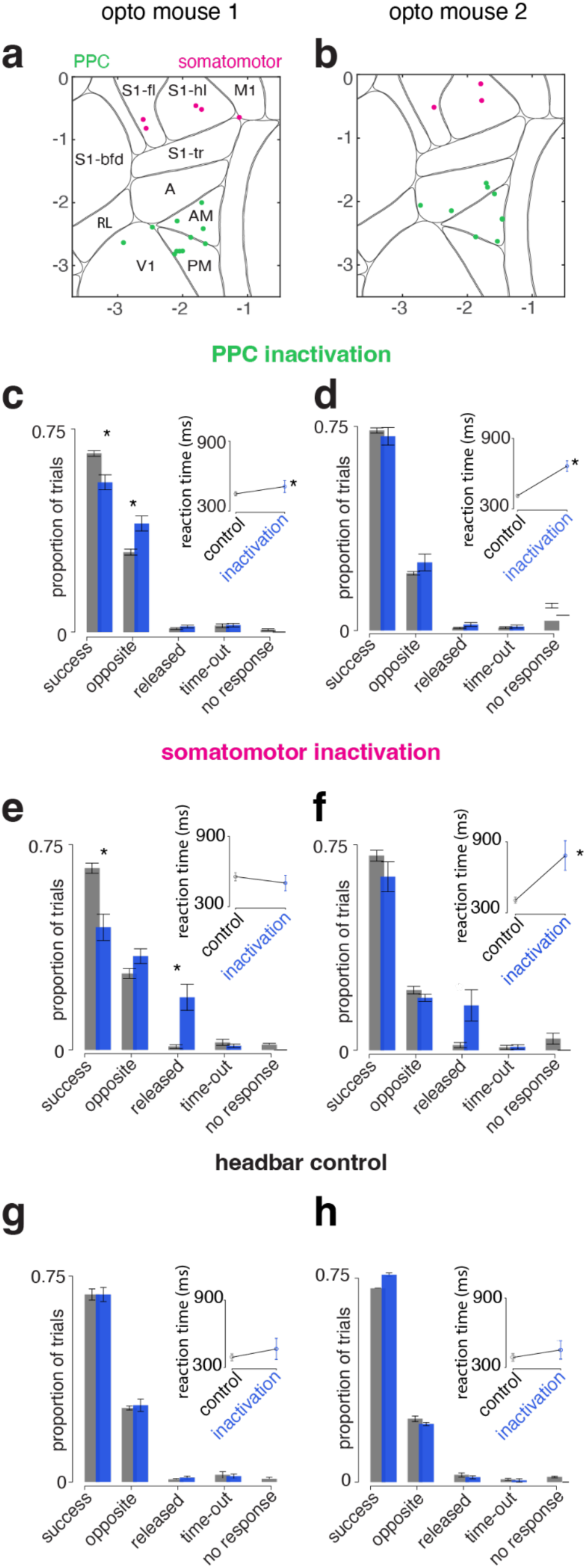
Effects of contralateral PPC, somatomotor inactivation and headbar control on behavior. (a) Centers of the different inactivation spots (sessions) from the first mouse superimposed on the Allen CCF. Green color indicates PPC inactivation sessions, pink color indicates somatomotor inactivation sessions. (b) Same as (a) for the second mouse. (c) Bar plot: average proportion of trials with various outcomes for control (unstimulated trials, gray) and contralateral PPC inactivation (blue) pooled across sessions in the first mouse. SEMs are across sessions (n=11 sessions). Stars indicate a difference at p=0.0025 and p=0.007 by Wilcoxon Rank Sum test for the outcome type “success” and “opposite” respectively. Inset shows the mean reaction times for control (n=2768) and inactivation trials (n=506). For reaction time analyses, only “clean success” trials are considered. Stars indicate a difference at p=0.035, t-test. (d) Same as (a) for second mouse (n=9 sessions). Inset shows the mean reaction times for control (n=2961) and inactivation trials (n=609). Stars indicate a difference at p=2×10^-12^, t-test. (e) Same bar plot as (a) for inactivation sessions over somatomotor areas (S1-fl, S1-hl, M1-fl) in mouse 1 (n=5 sessions). Stars indicate a difference at p=0.016 and p=0.008 by Wilcoxon Rank Sum test for the outcome type “success” and “release” respectively. Inset shows the mean reaction times for control (n=1143) and inactivation trials (n=159). (f) Second mouse (n=3 sessions). Inset shows the mean reaction times for control (n=968) and inactivation trials (n=153). Stars indicate a difference at p=4×10^-7^, t-test. (g) Bar plot: average proportion of trials with various outcomes for control (unstimulated trials, gray) and headbar control (blue, light ‘stimulation’ applied over the headbar) pooled across sessions in the first mouse. SEMs are across sessions (n=4 sessions). Inset shows the mean reaction times for control (n=937) and inactivation trials (n=186). (h) Same as (g) for second mouse (n=2 sessions). Inset shows the mean reaction times for control (n=665) and inactivation trials (n=162).

**Supplementary figure 7.**
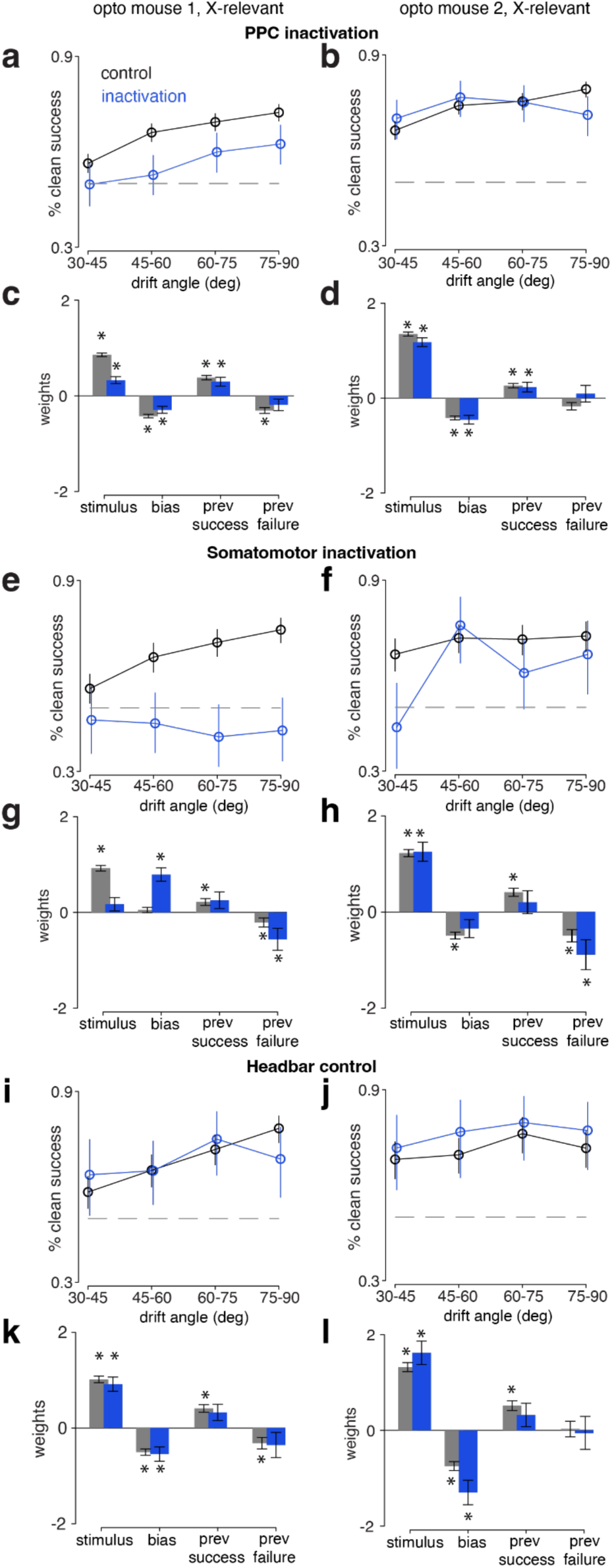
Quantifications of behavior and choice models for inactivation sessions. (a) Proportion of trials that were successful and had “clean” movements as a function of visual drift angle for control (gray) and inactivation (blue) trials in mouse 1. SEMs are across sessions; n=11 sessions, 4194 control and 897 inactivation trials over PPC. (b) Same as (a) for mouse 2; n=9 sessions, 4014 control and 831 inactivation trials over PPC. (c) Estimated weights from logistic regression choice model for the stimulus (signed relevant visual information) and strategy variables (bias, previous success and previous failure) for control (gray) and inactivation (blue) sessions. Control and inactivation trials were concatenated across different sessions and two global models were fit. SEMs obtained from the model fits. Stars indicate p<0.05, estimates obtained from the model fit. (d) Same as (c) for mouse 2. (e) Same plot as (a) for inactivation sessions over somatomotor areas (S1-fl, S1-hl, M1-fl) in mouse 1; n=5 sessions, 1709 control and 364 inactivation trials. (f) Same as (e) for mouse 2; n=3 sessions, 1367 control and 244 inactivation trials. (g) Same as (c) for mouse 1, inactivation sessions over SM. (h) Same as (g) for mouse 2. (i) Same plot as (a) for sessions with light ‘stimulation’ applied over the headbar as a control in mouse 1; n=4 sessions, 1369 control and 273 stimulation trials. (j) Same as (i) for mouse 2; n=2 sessions, 934 control and 212 stimulation trials. (k) Same as (c) for mouse 1, control sessions over the headbar. (l) Same as (k) for mouse 2.

**Supplementary figure 8:**
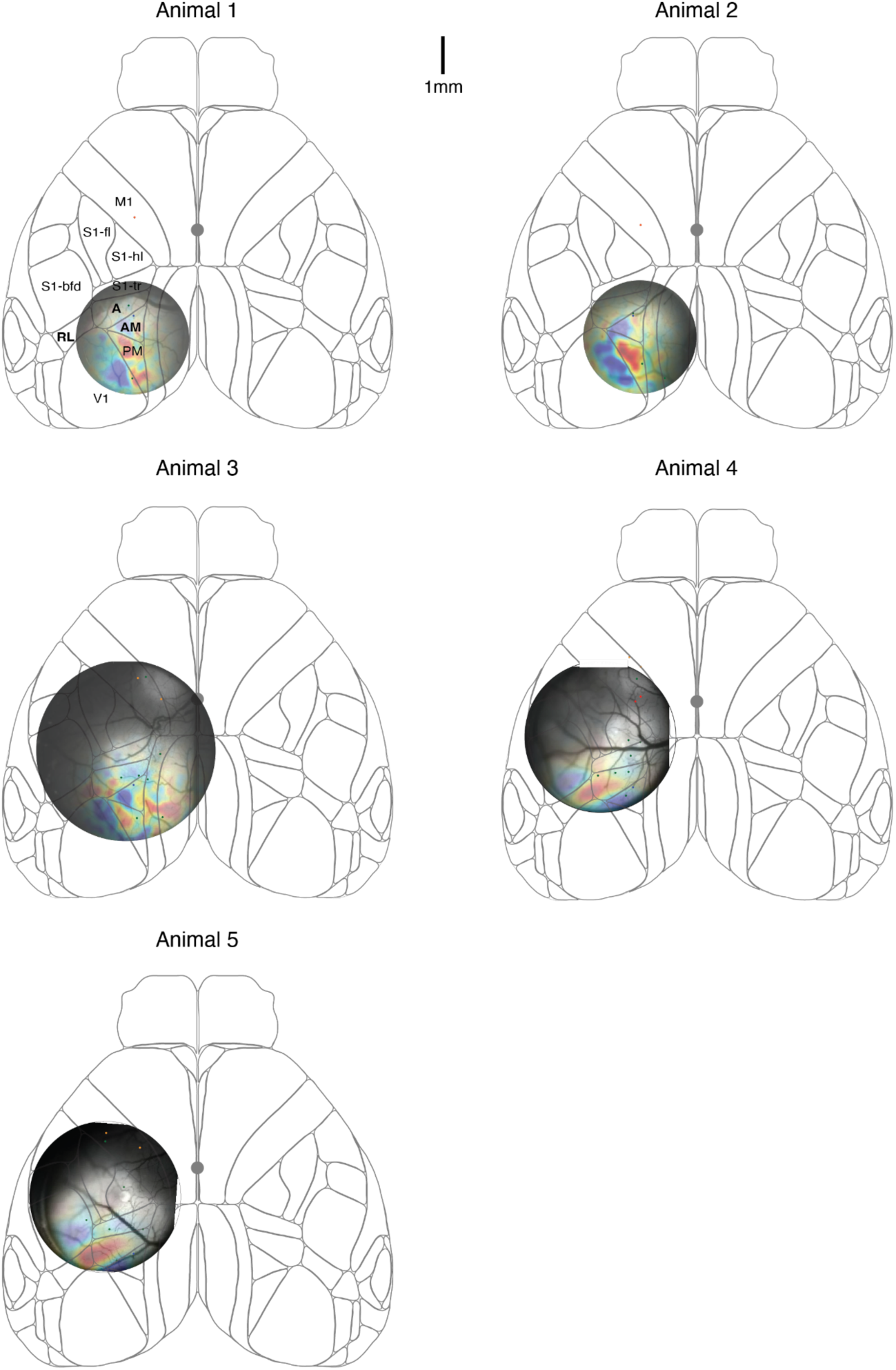
Retinotopic mapping to identify PPC. Widefield window images with superimposed retinotopic maps aligned to the Allen CCF for all imaging mice. Within each window, green dots are the AAV-Cre injection locations (to induce GCaMP6f expression) and orange dots are AAVretro-tdTomato injection locations (to label projection neurons). Scale bar indicates 1mm. Bold names indicate the areas that constitute PPC: A, AM, and RL.

**Supplementary figure 9.**
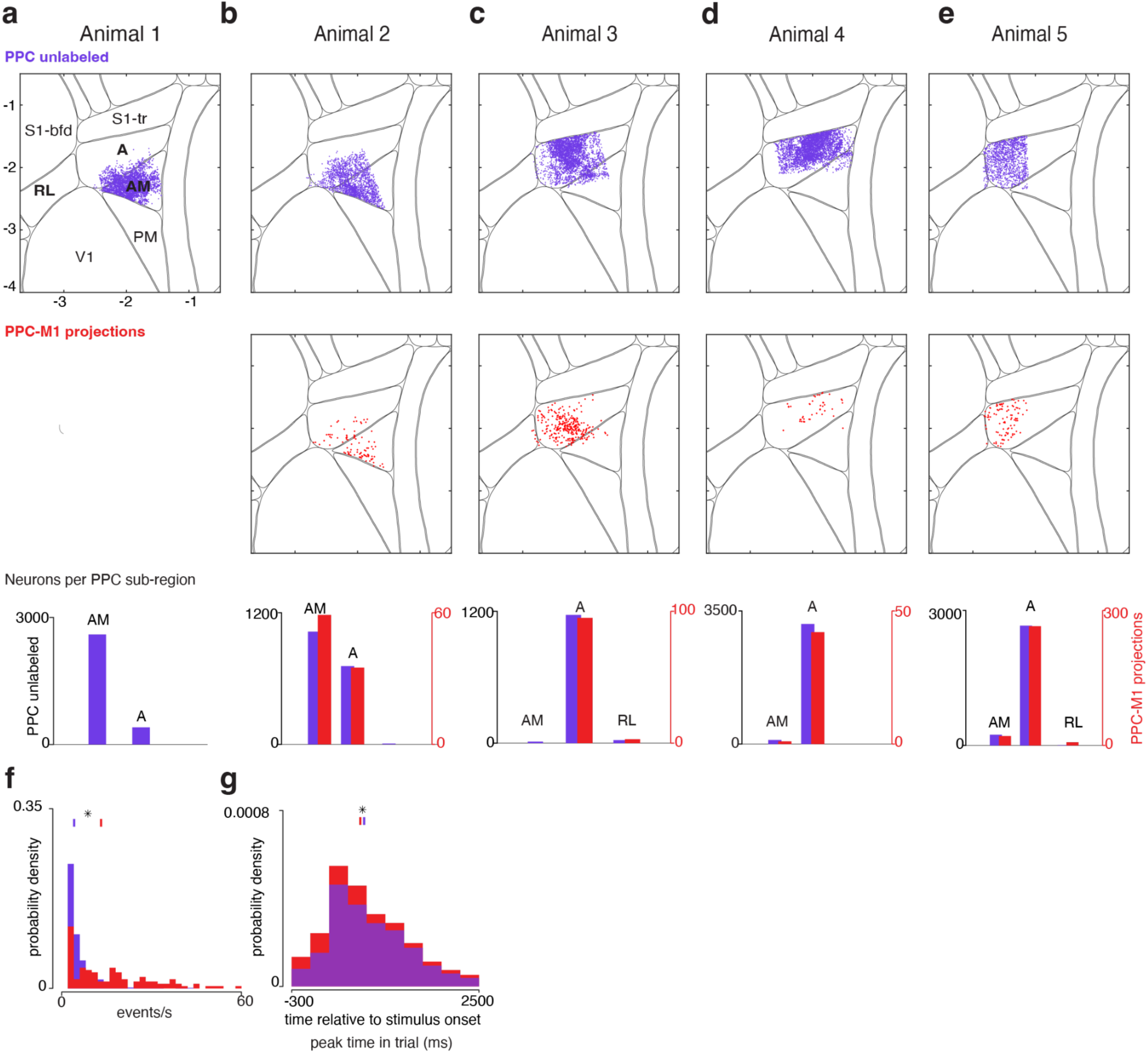
FOV locations and neuron counts per mouse. (a-e) Locations of imaged neurons (in PPC) with purple showing imaged cells without red (“unlabeled”) in each expert mouse used for imaging experiments (top row), locations of red-labeled M1 projection neurons in red (middle row) and the total number of neurons per sub-region of PPC (bottom row). Retrograde labeling was not successful in mouse 1, but “unlabeled” cells were included in analysis. (f) Histogram of firing rate distributions in unlabeled and PPC-M1 projection neurons. Lines denote medians (significant difference, Wilcoxon Rank Sum test, p=7×10^-19^). (g) Histogram of the time of peak activity during a trial. Distributions are truncated for visual clarity. Medians were significantly different (Wilcoxon Rank Sum test, p=6×10^-20^).

**Supplementary figure 10.**
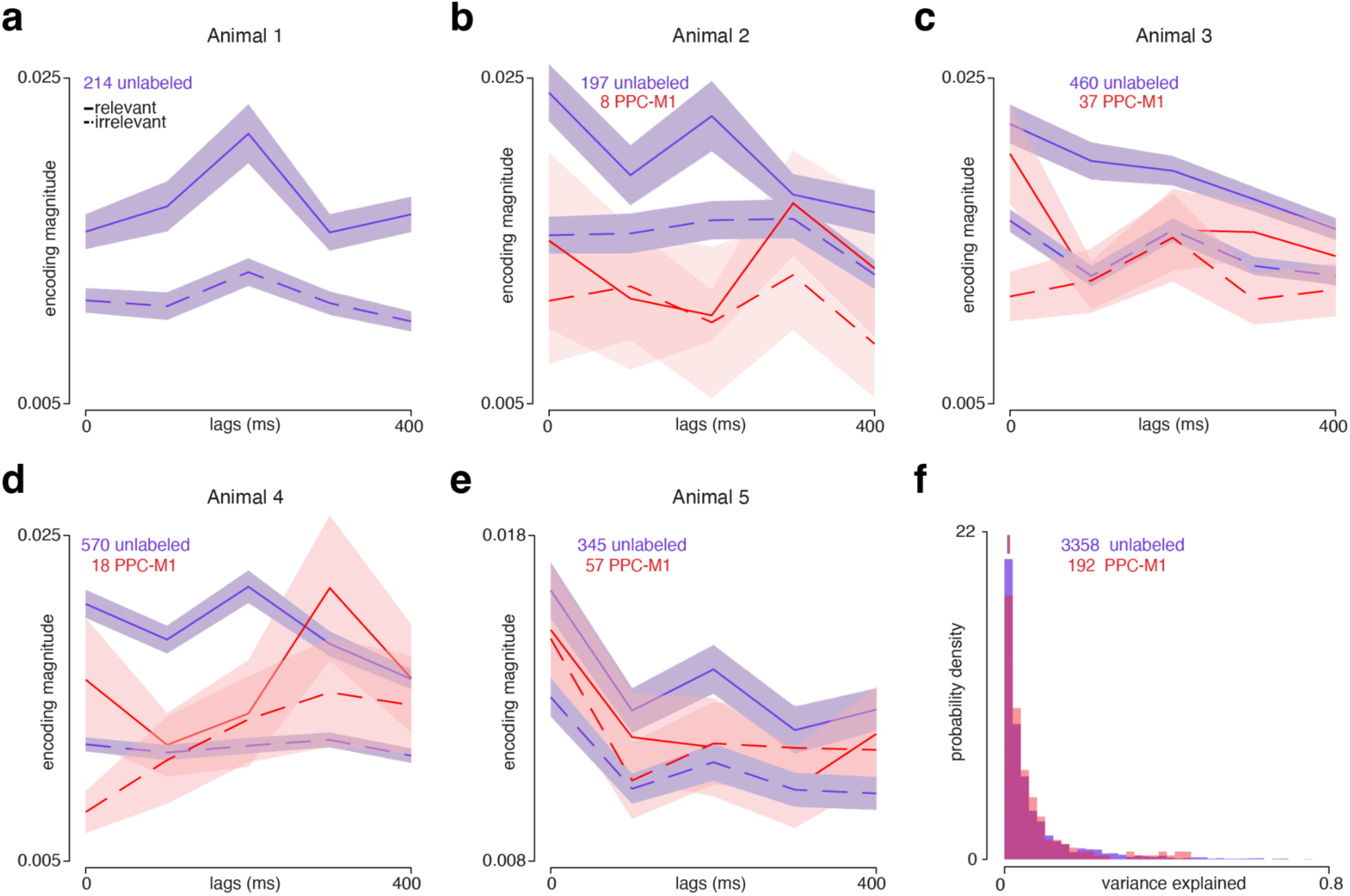
Visual encoding kernels for individual mice and quantification of random mixed selectivity. (a-e) Population averaged task-relevant and irrelevant visual encoding kernels for animals 1-5. Note that animal 1 did not have any projection neurons labeled. (f) Histogram of the single-trial variance explained by the encoding model for unlabeled (purple) and PPC-M1 (red) projection neurons.

**Supplementary figure 11.**
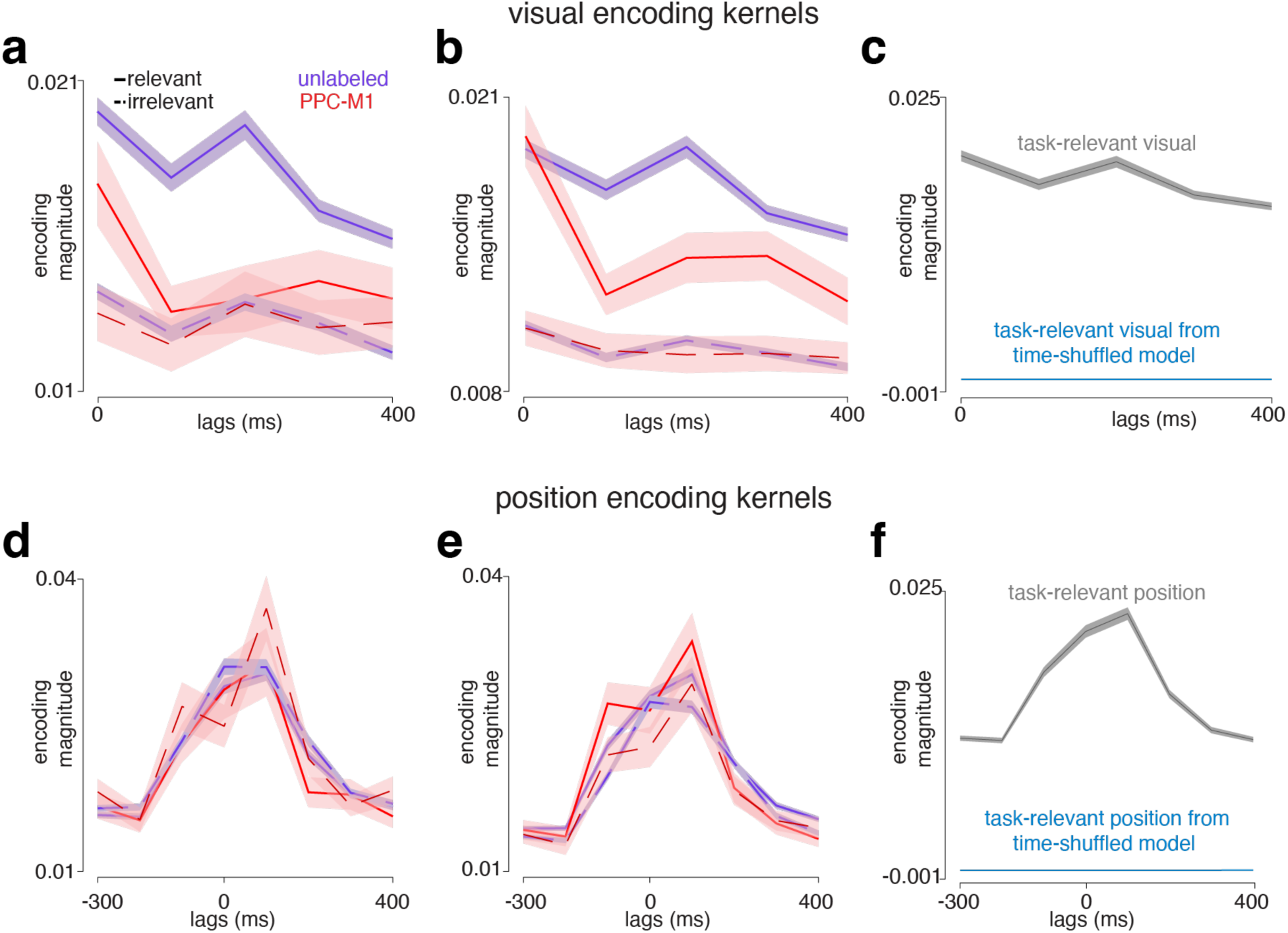
Encoding model controls. (a) Population-averaged task-relevant and irrelevant visual encoding kernels where relevant and irrelevant movement axes were as defined by the task instead of the principal components (PCs) of movement in the design matrix. Results were very similar to using the PCs. (b) Population-averaged task-relevant and irrelevant visual encoding kernels when all trials were used to fit the model. For Figures 4 and 5, only trials with equal relevant and irrelevant information in the visual stimulus were used for model fitting. (c) Population-averaged task-relevant visual encoding kernel across all well-fit neurons in PPC (gray) and the population averaged task-relevant visual encoding kernel computed from the time-shuffled model (blue). SEMs are across neurons. (d-f) Same as (a-c) for the position kernels.

